# Bacterial lipoate protein ligases rescue lipoylation and respiration deficiency in mammals

**DOI:** 10.1101/2024.11.22.624956

**Authors:** Zhijuan Hu, Junru Yu, Ziwei Liu, Min Jiang, An-Ping Zeng

**Affiliations:** Center of Synthetic Biology and Integrated Bioengineering, Westlake University, Hangzhou, Zhejiang, China; School of Engineering, Westlake University, Hangzhou, Zhejiang, China; School of Life Sciences, Westlake University, Hangzhou, Zhejiang, China; Research Center for Industries of the Future, Westlake University, Hangzhou, Zhejiang, China; Key Laboratory of Intelligent Low-Carbon Biosynthesis of Zhejiang Province, Westlake University, Hangzhou, Zhejiang, China

**Keywords:** Lipoylation deficiency, bacterial salvage lipoylation pathway, lipoate protein ligase, mitochondrial energy metabolism, gene therapy

## Abstract

Human lipoylation pathway deficiencies caused by LIPT2/LIAS/LIPT1 gene mutations lead to inherited metabolic disorders characterized by severe defects of mitochondrial energy and amino acids metabolism. Patients with such mutations suffer from hyperlactic acidemia, encephalopathy, hypotonia and early death. So far there is no effective treatment. Here we introduced the bacteria salvage lipoylation pathway, which is absent in eukaryotes, into lipoylation defective human cells and mouse models. Both *E. coli*-derived LplA and *B. subtilis*-derived LplJ restored the phenotypes of LIPT2/LIAS/LIPT1 gene knockout cells to those of wildtype cells with lipoic acid supplementation. Biochemical and isotope tracing analysis demonstrated LplA and LplJ function through direct lipoylation of the H protein of glycine cleavage system and the E2 subunits of 2-oxoacid dehydrogenases, thereby reactivating the short-circuited TCA cycle and amino acids metabolism. This strategy is further extended to treat lipoyl precursor supply defects caused by MECR and FDX1 mutations and its efficacy and safety are validated in *Lipt1^-/-^* and *LplA^OE/+^* mouse models. This study provides systematic characterization of lipoylation deficiency and paves the way to gene therapy for treatment of lipoylation-deficient patients.

## Introduction

Lipoylation is an essential posttranslational modification (PTM) for the function of key metabolic enzymes that involves the covalent attachment of a lipoyl group to a lysine residue of a target protein, providing a “swinging arm” to facilitate substrate channeling and electron transfer^1–4^. Lipoylation represents a rare yet conserved PTM across all organisms. To date, lipoylation has been identified as essential for the function of five enzyme complexes: the glycine cleavage system (GCS) and four 2-oxoacid dehydrogenases, namely, pyruvate dehydrogenase (PDH), 2-oxoglutarate dehydrogenase (OGDH), branched chain 2-oxoacid dehydrogenase (BCKDH), and 2-oxoadipate dehydrogenase (OADH)^5,6^. These enzyme complexes share a similar architecture with multiple subunits. The lipoyl group is bound to the E2 subunit of the four 2-oxoacid dehydrogenases and to the H protein of GCS. Additionally, lipoylation of the E3-binding protein (PDHX) of PDH has been reported^7^. In eukaryotes, these lipoylation-dependent proteins (LDP) are exclusively located in mitochondria. PDH and OGDH are involved in the TCA cycle, while BCKDH, OADH and GCS are engaged in the catabolism of branched-chain amino acids (valine, leucine, isoleucine), lysine and glycine, respectively. Consequently, lipoylation is crucial for both mitochondrial energy metabolism and amino acid metabolism.

The lipoylation biosynthetic pathways in bacteria, yeast, and humans have been extensively characterized^8–11^. There are two types of distinctive pathways: *de novo* synthesis and salvage pathways. According to current knowledge, the *de novo* lipoylation pathway in humans is catalyzed sequentially by three enzymes (**Figure 1a**): (1) octanoyltransferase (LIPT2) transfers the octanoyl group from octanoyl-ACP, produced via mitochondrial fatty acid synthesis, onto the H protein; (2) lipoyl synthase (LIAS) subsequently inserts two sulfur atoms from iron-sulfur cluster at the C6 and C8 positions of the octanoyl group to form lipoyl-H; (3) lipoyltransferase (LIPT1) then transfers the lipoyl group to the E2 subunits of the other four 2-oxoacid dehydrogenases. The bacterial *de novo* pathway in *Bacillus subtilis* is similar to that in humans but differs in *E. coli*, where LipB directly transfers octanoyl group to H protein and E2 subunits, followed by LipA-catalyzed sulfur insertion to form lipoyl group (Figure 1a). Notably, an additional salvage lipoylation pathway has been reported in several bacteria, including *E. coli*, *B. subtilis*, and *Streptomyces coelicolor*^8,12,13^. A lipoate protein ligase, like LplA from *E. coli* or LplJ from *B. subtilis*, activates exogenous lipoic acid through adenylation with ATP and transfers lipoyl group from lipoyl-AMP to H protein and E2 subunits. In contrast, humans lack LplA homologues and cannot utilize exogenous lipoic acid for protein lipoylation. Although an acetyl-CoA synthetase medium-chain family member 1 (ACSM1) has been reported to activate lipoic acid with GTP, there is no substantial evidence to prove that ACSM1 contributes to protein lipoylation *in vivo*^14,15^.

**Figure 1.**
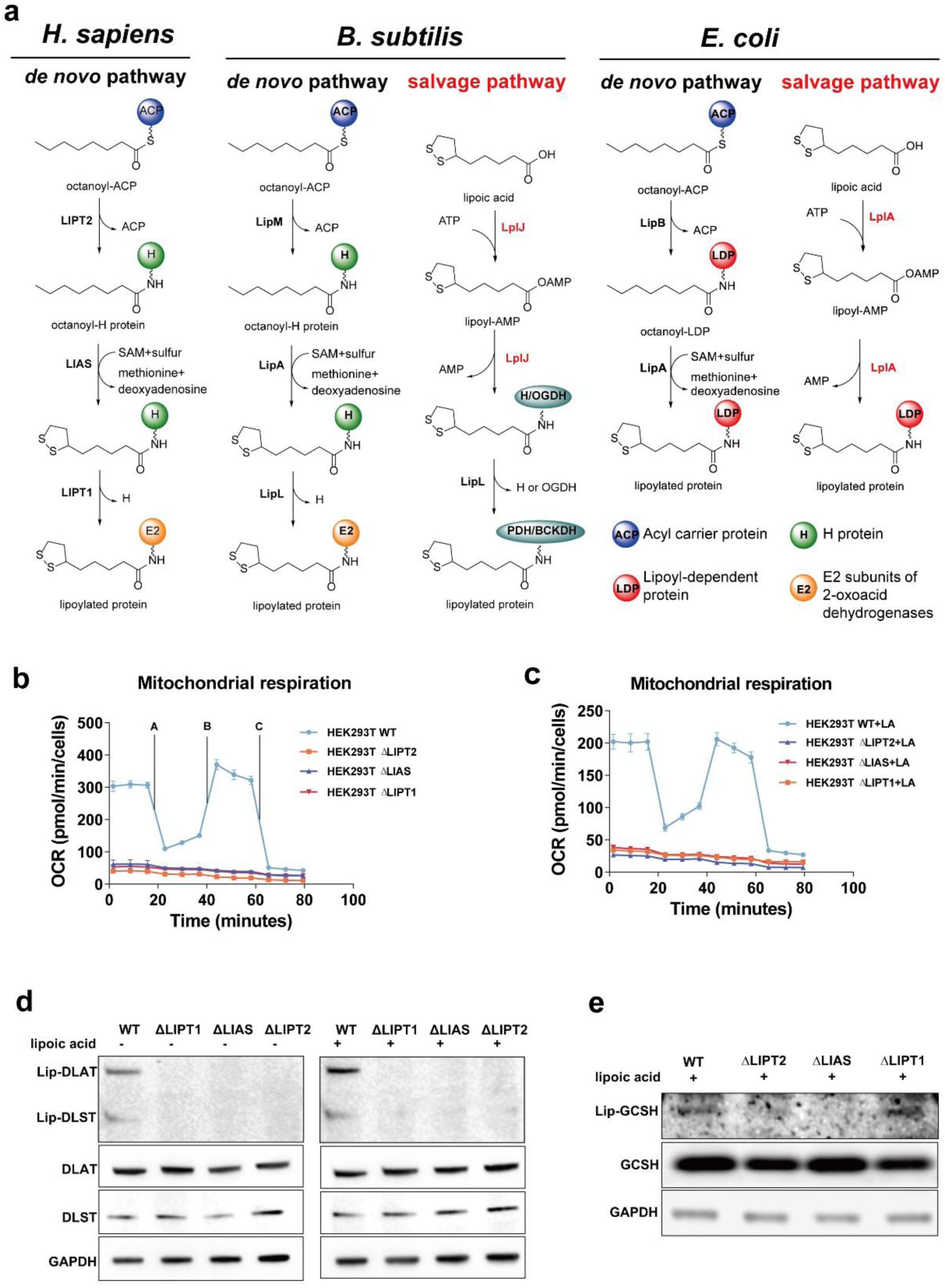
Lipoylation pathways, severe mitochondrial respiration defect, protein lipoylation deficiency and growth arrest in human HEK293T cells caused by disruption of lipoylation. (a) Lipoylation pathways in *H. sapiens*, *B. subtilis*, and *E. coli*. (b) Mitochondrial respiration of HEK293T WT, ΔLIPT2, ΔLIAS, and ΔLIPT1 cells without lipoic acid supplementation as determined by Seahorse Mito Stress Test. OCR were measured through sequential injection of drug A (2 μM oligomycin), drug B (0.5 μM carbonyl cyanide-4 (trifluoromethoxy) phenylhydrazone (FCCP)), drug C (1 μM rotenone and 1 μM antimycin A). (c) Mitochondrial respiration of HEK293T WT, ΔLIPT2, ΔLIAS, and ΔLIPT1 cells with lipoic acid supplementation as determined by Seahorse Mito Stress Test. (d) Lipoylation levels of DLAT and DLST of HEK293T WT, ΔLIPT2, ΔLIAS, and ΔLIPT1 cells with presence or absence of lipoic acid as determined by Western blotting analysis, GAPDH, total DLAT, and total DLST were used as controls. (e) Lipoylation level of GCSH of HEK293T WT, ΔLIPT2, ΔLIAS, and ΔLIPT1 cells with presence or absence of lipoic acid as determined by Western blotting analysis, GAPDH and total GCSH were used as controls.

Mutations in genes involved in lipoylation biosynthesis have been associated with mitochondrial disorders characterized by severe defects in energy metabolism^2,3^. LIAS (OMIM#607031) mutations have been identified in five patients, all presenting with hyperglycinemia, lactic acidosis, and seizures^16–19^. Typical biochemical phenotypes include elevated glycine levels in both serum and cerebrospinal fluid, decreased or absent protein lipoylation, and deficient activities of GCS, PDH, and OGDH enzymes. LIPT2 (OMIM#617659) mutations have been reported in three patients with severe neonatal encephalopathy, lactic acidosis, and brain abnormalities^20^. These patients exhibited moderately increased plasma glycine levels. Fibroblasts from these patients showed reduced mitochondrial respiration and decreased protein lipoylation. LIPT1 (OMIM#610284) deficiencies have been documented in six patients with fatal lactic acidosis, psychomotor regression, Leigh syndrome, and epileptic encephalopathy^21–25^. Biochemical profiles revealed elevated serum and urine lactate, elevated urine ketoglutarate, and a specific lipoylation defect of the 2-oxoacid dehydrogenases. Unlike LIAS or LIPT2 mutations, glycine levels remained normal in LIPT1 mutations, consistent with its role in lipoyl transfer from lipoyl-H to E2 subunits, which distinguishes the diagnosis of LIPT1 from the other two mutations. Besides the genes involved in lipoylation biosynthesis, those involved in mitochondrial fatty acid synthesis (ACP, MECR)^26,27^, iron-sulfur cluster synthesis (NFU1, BOLA3, IBA57, GLRX5)^18^, and lipoyl group recycling (DLD)^28^ also exhibited compromised protein lipoylation and LIAS-like phenotypes.

Currently, there is no effective treatment for alleviating the suffering of these patients. Oral administration of lipoic acid or extracellular supplementation of lipoic acid in patients’ fibroblasts does not rescue the lipoylation deficiency. A recent study identified a cocktail of antioxidants, including pantothenate, nicotinamide, vitamin E, thiamine, biotin, and α-lipoic acid, which improved the phenotypes of a LIPT1 mutation in patient-derived cell models^29^. Introducing a bacterial lipoate ligase into the mitochondria has been proposed in a previous review^4^. Indeed, the complementation of mitochondrially localized LplA with lipoic acid demonstrated a rescue effect in Δlip2 (LIPT2 homolog), Δlip5 (LIAS), and Δgcv3 (GCSH) mutant yeast strains, but not in the Δlip3 (LIPT1) strain, which is due to the successful lipoylation of the yeast H protein and the E2 subunit of PDH by LplA, but not the E2 subunit of OGDH^30^.

Here we report the introduction of bacterial lipoate ligases into human lipoylation deficient cell models. With lipoic acid supplementation, both LplA and LplJ restored the deficient mitochondrial energy metabolism of ΔLIPT2, ΔLIAS, and ΔLIPT1 cells to levels comparable to wild-type cells. We proved that LplA and LplJ function through direct lipoylation of the E2 subunits of PDH and OGDH, reactivating the short-circuited TCA cycle by using *in vitro* and *in vivo* protein lipoylation level measurement, PDH and OGDH activity detection, enzyme mutagenesis study, and isotope-labeled carbon tracing. This treatment strategy was further expanded to lipoyl precursor supply (MECR) and sulfur insertion accessary partner (FDX1) defects but not to lipoyl recycling (DLD) defects. Finally, we evaluated the efficacy and safety of this strategy in *Lipt1* knockout and *LplA* knockin mice models. Interbreeding between heterozygous *Lipt1*^+/-^ *LplA*^OE/+^ mice with oral administration of lipoic acid successfully gave birth to homozygous *Lipt1*^-/-^ mice carrying either heterozygous *LplA*^OE/+^ alleles or homozygous *LplA*^OE/OE^ alleles, rescuing embryonic demise caused by homozygous *Lipt1*^-/-^ mutations. The body composition, energy expenditure and hematology of *LplA^OE/+^* mice exhibited normal to wildtype mice, indicating the safety of introducing LplA *in vivo*.

## Results and Discussion

### Lipoylation pathway disruption causes severe mitochondrial respiration defect, protein lipoylation deficiency and growth arrest in human cells

To unravel the potential role of bacterial lipoate protein ligases in human lipoylation deficiency, we first generated LIPT2, LIAS, and LIPT1 knockout human embryonic kidney (HEK293T) cells using CRISPR-Cas9-mediated gene editing technology (**Figure S1a-d**). As mitochondrial energy metabolism is a characteristic feature of lipoylation deficiency, we examined mitochondria-related parameters in the knockout cells. Compared to wildtype controls, HEK293T ΔLIPT2, ΔLIAS, and ΔLIPT1 cells all exhibited markedly suppressed mitochondrial respiration, as evidenced by significantly impaired basal and maximal respiration as well as decreased ATP production (**Figure 1b**). Of note, the addition of 20 μM lipoic acid to the culture medium failed to improve mitochondrial respiration (**Figure 1c**), consistent with previous studies^20^. Subsequently, the loss of protein lipoylation of DLAT and DLST, the E2 subunits of PDH and OGDH respectively, was confirmed by Western blotting analysis (**Figure 1d**). Consistently, the presence or absence of lipoic acid had no influence on protein lipoylation level. Since the lipoylation of the H protein differentiates the defect of LIPT1 from LIPT2 and LIAS, we also assessed GCSH lipoylation with particular attention in the knockout cells. The GCSH lipoylation level in ΔLIPT1 cells was identical to that in wild-type cells, but not in Δ LIPT2 and Δ LIAS cells (**Figure 1e**). During cell cultivation, we observed a marked cell proliferation arrest in the knockout cells, and the media rapidly turned yellow. The cell growth curve assay confirmed that the growth of the deficient cells was arrested compared to the wild-type cells (**Figure S1e**). In addition, given that intracellular metabolism is another feature associated with lipoylation, we employed untargeted metabolomics profiling of these cells. We found that 2-oxoglutaric acid, 2-oxoisocaproic acid, and 2-oxoadipic acid, the substrates of OGDH, BCKDH, and ADH respectively, were significantly accumulated in knockout cells while fumaric acid and malate, the intermediates of TCA cycle were significantly reduced, indicating impaired lipoylated proteins (**Figure S2**). Altogether, our data suggest that LIPT2, LIAS, and LIPT1 deficiency in HEK293T cells induce typical mitochondrial energy metabolism defects, protein lipoylation deficiency and growth arrest, resembling the phenotypes of patient fibroblasts, making them suitable models for subsequent functional studies.

### LplA and LplJ rescue lipoylation deficiency in HEK293T cells

We then sought to investigate the rescue potential of bacterial lipoate ligases against human lipoylation defects. Specifically, LplA derived from *E. coli* and LplJ from *B. subtilis* were assembled with a mitochondrial targeting signal (MTS) and a GFP tag to generate MTS-GFP-LplA/LplJ cassettes, facilitating mitochondrial localization and cell monitoring (**Figure 2a**). Catalytic activity of LplA was retained in GFP-LplA fusion protein, as evidenced by *in vitro* biochemical assays (**Figure S3a-b**). The MTS-GFP-LplA/LplJ constructs as well as MTS-GFP control were stably expressed by lentiviral transduction of HEK293T ΔLIPT2, ΔLIAS, and ΔLIPT1 cells respectively. Their cellular localization was confirmed by immunofluorescence analysis that all the constructs with MTS were exclusively localized inside the mitochondria as supported by the overlap of GFP signals with the mitochondrial marker tom20 (**Figure S3d-e**).

**Figure 2.**
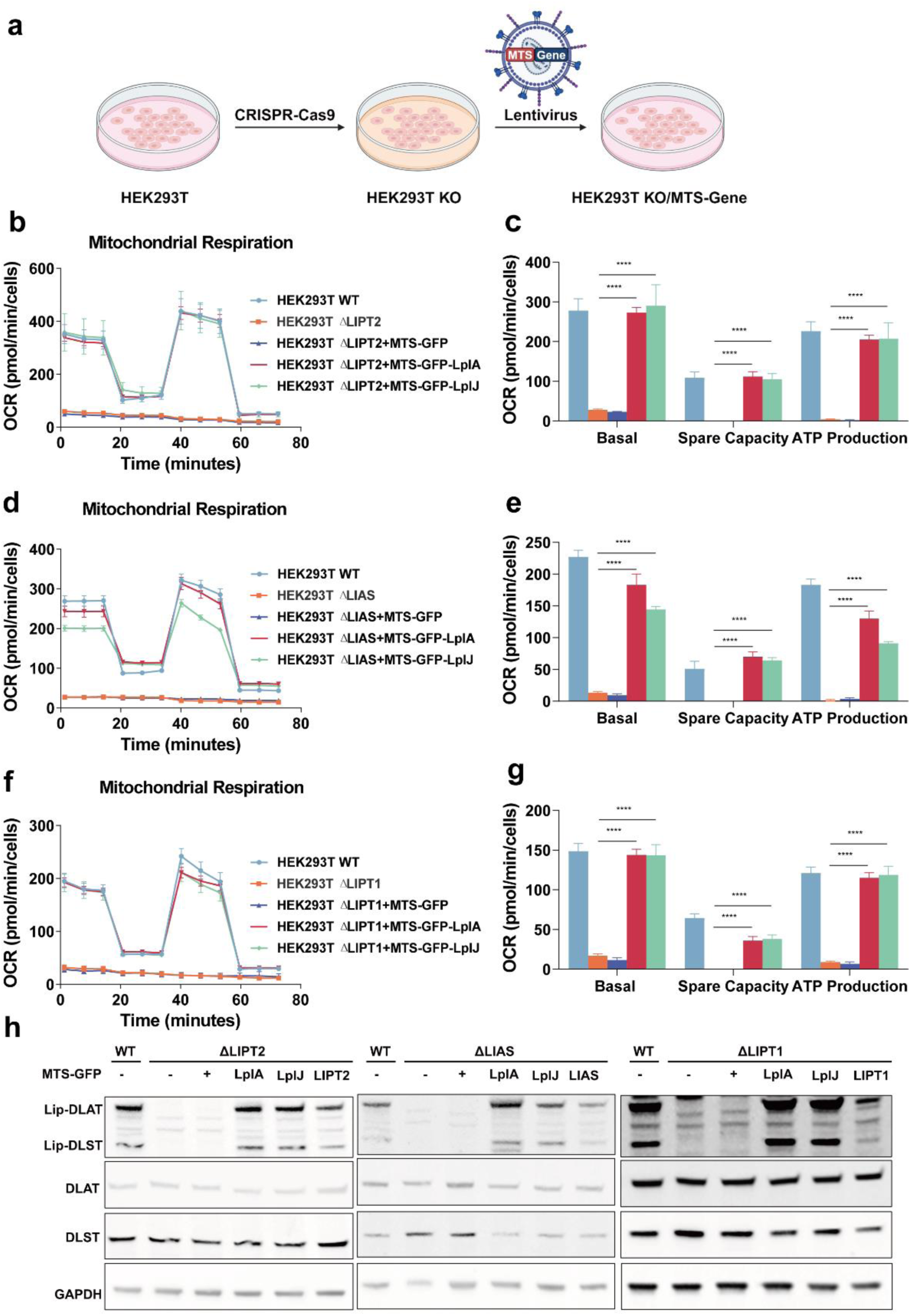
LplA and LplJ rescue lipoylation deficiency with lipoic acid supplementation in HEK293T cells. (a) Scheme of generating HEK293T knockout cells and complementary cells. (b) Mitochondrial respiration of HEK293T WT, ΔLIPT2 and related complementary cells with lipoic acid supplementation. (c) OCR-related basal respiration, spare capacity and ATP production of ΔLIPT2 related cells. (d) Mitochondrial respiration of HEK293T WT, ΔLIAS and related complementary cells with lipoic acid supplementation. (e) OCR-related basal respiration, spare capacity and ATP production of ΔLIAS related cells. (f) Mitochondrial respiration of HEK293T WT, ΔLIPT1 and related complementary cells with lipoic acid supplementation. (g) OCR-related basal respiration, spare capacity and ATP production of ΔLIPT1 related cells. (h) Lipoylation levels of DLAT and DLST of related cells were determined by Western blotting, GAPDH, total DLAT, and total DLST were used as controls. Data represented as mean±SD. Statistical significance was determined using multiple t tests (\**p* <0.05, \*\**p* <0.01, \*\*\**p* <0.001, and \*\*\*\**p* <0.0001).

We then examined the mitochondrial energy metabolism in the supplementary cells along with knockout cells and wildtype cells. With addition of lipoic acid, both MTS-GFP-LplA and MTS-GFP-LplJ restored mitochondrial respiration capacity to nearly wild-type levels, whereas the MTS-GFP control remained at the knockout cell level, manifested by the significantly enhanced basal respiration, spare respiration capacity and ATP production (**Figure 2b-g**). The rescue effect was consistently observed in all three knockout cell models. Concurrently, we detected resumed bands corresponding to lip-DLAT and lip-DLST upon the expression of LplA or LplJ (**Figure 2h**). Without addition of lipoic acid, the rescue effect was abolished in LIPT2 knockout cells but was partially retained in LIPT1 knockout cells (**Figure 3a-f**). We hypothesized that LIPT1 knockout would not affect the lipoylation of the H protein, and a small amount of free lipoic acid might be generated by endogenous lipoamidase Sirtuin 4 through the hydrolysis of lip-H^7^. In support of this, we observed partial recovery of DLAT lipoylation but not DLST lipoylation in LIPT1 knockout cells (**Figure 3g**). It is of note that intracellular lipoic acid was not detected by triple quadrupole mass spectrometry (data not shown).

**Figure 3.**
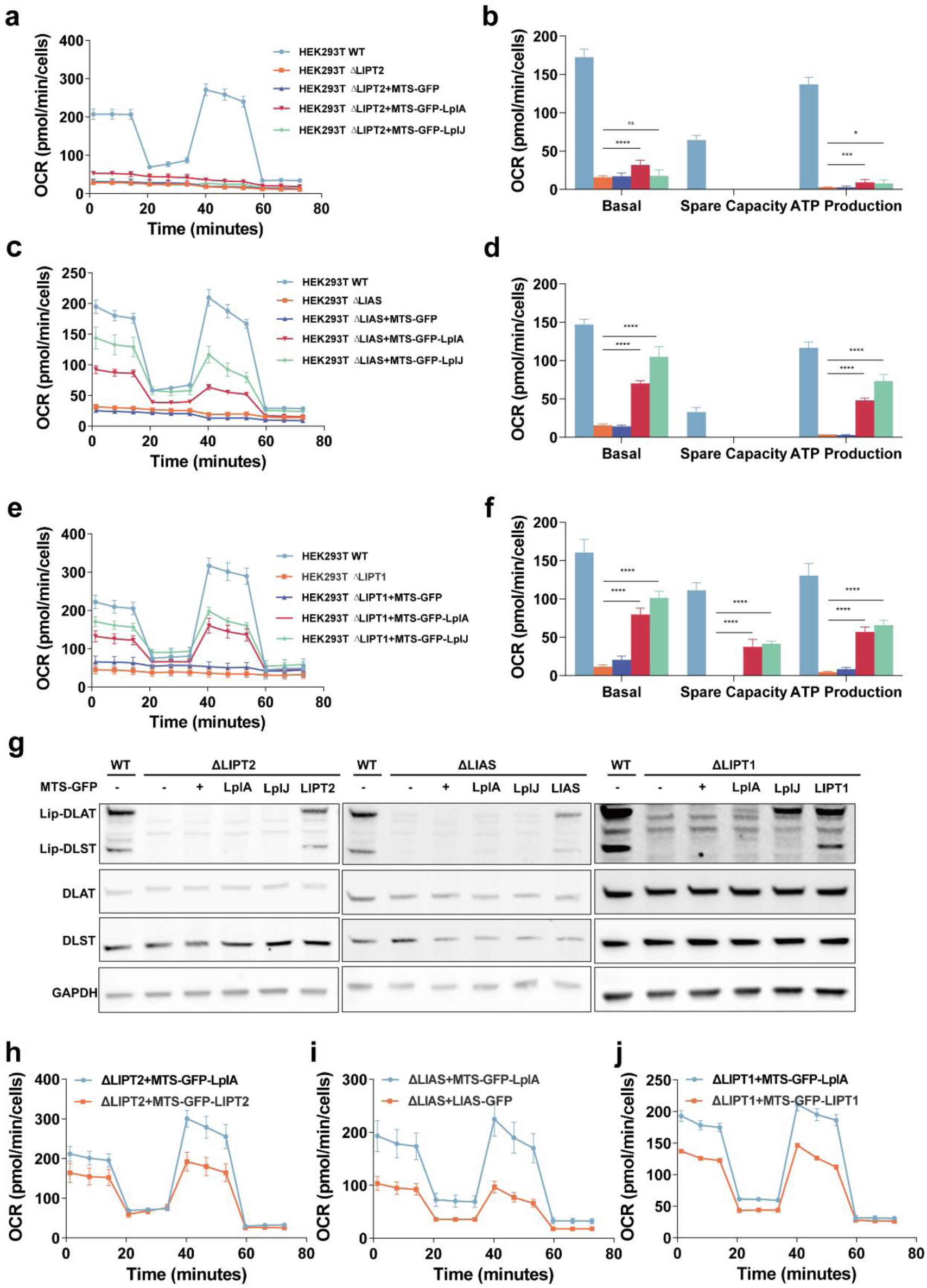
LplA and LplJ attenuate rescue effect without lipoic acid supplementation. (a) Mitochondrial respiration of HEK293T WT, Δ LIPT2 and related complementary cells without lipoic acid supplementation. (b) OCR-related basal respiration, spare capacity and ATP production of ΔLIPT2 related cells. (c) Mitochondrial respiration of HEK293T WT, ΔLIAS and related complementary cells without lipoic acid supplementation. (d) OCR-related basal respiration, spare capacity and ATP production of ΔLIAS related cells. (e) Mitochondrial respiration of HEK293T WT, Δ LIPT1 and related complementary cells without lipoic acid supplementation. (f) OCR-related basal respiration, spare capacity and ATP production of ΔLIPT1 related cells. (g) Lipoylation levels of DLAT and DLST of related cells without lipoic acid were determined by Western blotting, GAPDH, total DLAT, and total DLST were used as controls. (h) Mitochondrial respiration of HEK293T ΔLIPT2 cells complemented with MTS-GFP-LplA or MTS-GFP-LIPT2 with lipoic acid supplementation. (i) Mitochondrial respiration of HEK293T ΔLIAS cells complemented with MTS-GFP-LplA or LIAS-GFP with lipoic acid supplementation. (j) Mitochondrial respiration of HEK293T ΔLIPT1 cells complemented with MTS-GFP-LplA or MTS-GFP-LIPT1 with lipoic acid supplementation. Data represented as mean±SD. Statistical significance was determined using multiple t tests (\**p* <0.05, \*\**p* <0.01, \*\*\**p* <0.001, and \*\*\*\**p* <0.0001).

To further evaluate the therapeutic potential of bacterial lipoate ligases in the treatment of human lipoylation deficiencies, we compared their effects with those of native gene supplements. LIPT2, LIAS, and LIPT1 were introduced into the cells in the same manner as LplA. Mitochondrial respiration was assessed in knockout cells complemented with the corresponding native genes in the presence of lipoic acid. The results indicated that LplA performed better than LIPT2, LIAS, and LIPT1 (**Figure 3h–3j**). We speculated that the MTS and GFP tag might influence the catalytic function of LIPT2, LIAS, and LIPT1. Indeed, MTS-GFP-LIAS showed no detectable improvement in LIAS knockout cells, prompting us to construct LIAS-GFP for comparison. A more effective complementation strategy for LIPT2, LIAS, and LIPT1 requires further investigation. Collectively, our findings demonstrate that LplA and LplJ can rescue lipoylation deficiencies in HEK293T cells and restore phenotypes to wild-type levels with lipoic acid supplementation.

### LplA and LplJ function through direct lipoylation on human proteins

To elucidate the mechanism by which bacterial lipoate ligases rescue human lipoylation deficiencies, we first investigated the molecular basis of the lipoylation reactions *in vitro* and *in vivo*. Human-derived DLAT, DLST and GCSH proteins, with their mitochondrial targeting sequences removed, were overexpressed and purified from *E. coli*. *In vitro* biochemical assays, conducted with and without LplA, demonstrated that LplA can lipoylate human E2 subunits and the H protein, as confirmed by western blotting analysis using anti-His tag and anti-lipoic acid antibodies to detect total and lipoylated proteins, respectively (**Figure 4a-c**). Notably, LplJ was found to lipoylate human DLAT, DLST, and GCSH, despite previous reports indicating its activity was limited to the E2 subunit of OGDH and the H protein in *B. subtilis*^31^. Amino acids alignment revealed the conservation of the glutamate residue in human E2 subunits located three residues before the catalytic lysine residue(**Figure S3f**), which was reported as a recognition signal for the lipoylation reaction^32^. Subsequently, the enzyme activities of intracellular PDH and OGDH were determined by colorimetric assays. Results showed a significant reduction in LIPT1 knockout cells and a rejuvenation in LplA or LplJ complementary cells (**Figure 4d-e**).

**Figure 4.**
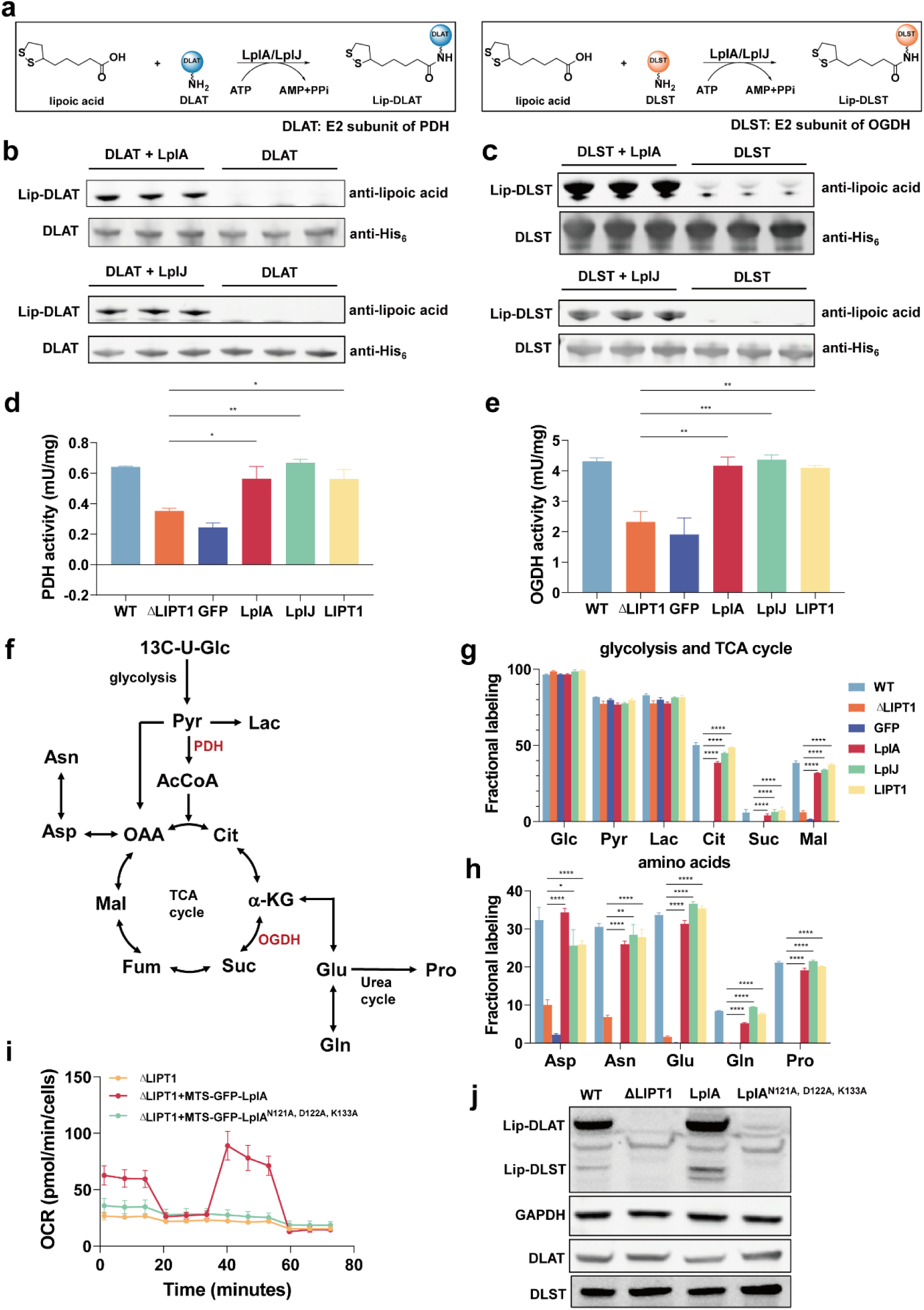
LplA and LplJ function through direct lipoylation on human proteins. (a) Schemes of lipoylation reactions of DLAT and DLST catalyzed by LplA or LplJ. (b) Lipoylation of human-derived DLAT by LplA or LplJ as determined by Western blotting analysis using anti-lipoic acid and anti-His tag antibodies. (c) Lipoylation of human-derived DLST by LplA or LplJ as determined by Western blotting analysis using anti-lipoic acid and anti-His tag antibodies. (d) PDH activities of HEK293T WT, ΔLIPT1, and ΔLIPT1 complemented with MTS-GFP, MTS-GFP-LplA, MTS-GFP-LplJ and MTS-GFP-LIPT1 cells with lipoic acid supplementation as determined by PDH activity colorimetric assay Kit. (e) OGDH activities of HEK293T WT, ΔLIPT1, and ΔLIPT1 complemented with MTS-GFP, MTS-GFP-LplA, MTS-GFP-LplJ and MTS-GFP-LIPT1 cells with lipoic acid supplementation as determined by OGDH activity colorimetric assay Kit. (f) Metabolic pathway of glycolysis, TCA cycle and closely related amino acids during 13C tracing analysis. (g) Fractional labeling of glycolysis and TCA cycle intermediates of HEK293T WT, Δ LIPT1, and ΔLIPT1 complemented with MTS-GFP, MTS-GFP-LplA, MTS-GFP-LplJ and MTS-GFP-LIPT1 cells after U-13C-glucose feeding. (h) Fractional labeling of amino acids. (i) Mitochondrial respiration of HEK293T ΔLIPT1, and ΔLIPT1 complemented with MTS-GFP-LplA and MTS-GFP-LplA^N121A, D122A, K133A^ cells with lipoic acid supplementation. (j) Lipoylation levels of DLAT and DLST of HEK293T ΔLIPT1, and ΔLIPT1 complemented with MTS-GFP-LplA and MTS-GFP-LplA^N121A, D122A, K133A^ cells with lipoic acid determined by Western blotting, GAPDH, total DLAT, and total DLST were used as controls. Data represented as mean±SD. Statistical significance was determined using multiple t tests (\**p* <0.05, \*\**p* <0.01, \*\*\**p* <0.001, and \*\*\*\**p* <0.0001).

Increases in PDH and OGDH activities predict enhanced oxidative metabolism. We then performed [U-^13^C] glucose feeding in wildtype, knockout and complementary cells (**Figure 4f-h & Figure S4**). Glycolysis intermediates displayed uniformly high fractional labeling rates across all cell lines whereas TCA cycle intermediates, specifically, citrate, succinate, and malate, showed dramatically low fractional labeling rates in knockout cells, indicating a short-circuited TCA cycle (**Figure 4g**). As expected, LplA and LplJ reactivated the TCA cycle as shown by the increased fractional labeling rates of TCA cycle intermediates (**Figure 4g**). In addition, we monitored the same pattern of fluctuations in the fractional labeling rates of aspartate, asparagine, glutamate, glutamine, and proline, which are closely exchanged with TCA cycle intermediates (**Figure 4h**). To obtain further evidence for the mode-of-action of LplA in regulating cellular energy metabolism, we generated an LplA mutant, LplA^N121A, D122A, K133A^, which abolishes its lipoylation activity (**Figure S3c**). Complementary experiments with this mutant confirmed that LplA^N121A, D122A, K133A^ indeed abrogates the recovery effect on mitochondrial respiration and protein lipoylation (**Figure 4i-j**). Altogether, our data proved that LplA and LplJ function through direct lipoylation on human lipoyl-dependent proteins, restored the function of PDH and OGDH, and reactivated mitochondrial energy metabolism and amino acids metabolism.

### Expanding the treatment scope to lipoyl precursor supply

The robust rescue effect of the bacterial salvage pathway in treating human lipoylation pathway defects has prompted us to expand its therapeutic scope. Mitochondrial fatty acid biosynthesis and iron-sulfur cluster synthesis are closely linked to lipoylation assembly (**Figure 5a**), providing octanoyl-ACP and sulfur atoms as essential building blocks, respectively^3^. Defects in related genes like MECR, ACP, NFU1, and BOLA3 are associated with human mitochondrial disorders^33^. Ferredoxin1 (FDX1) has been reported to be essential for protein lipoylation through direct binding to LIAS, thereby promoting its interaction with GCSH^34,35^. Dihydrolipoamide dehydrogenase (DLD) is the E3 subunit shared by all lipoyl-dependent enzymes, catalyzing the oxidation of dihydrolipoamide to form lipoamide^28^. We further applied LplA/LplJ complementation along with lipoic acid supplementation in HEK293T Δ MECR, Δ FDX1, and Δ DLD cells. Mitochondrial respiration was significantly enhanced upon expression of LplA or LplJ in ΔMECR and Δ FDX1 cells (**Figure 5b-e**), although not to wildtype levels, which is reasonable due to the additional roles of MECR in fatty acid synthesis and FDX1 in steroidogenesis^5,36^. In contrast, LplA or LplJ could not rescue basal, maximal, and ATP production-related respiration in ΔDLD cells (**Figure 5f-g**). Concurrently, protein lipoylation levels of DLAT and DLST were recovered in ΔMECR and ΔFDX1 cells, while ΔDLD cells retained partial protein lipoylation, which improved with LplA or LplJ expression (**Figure 5h**). We speculate that DLD deficiency does not affect the initial installation of the lipoyl group but restricts the extent of protein lipoylation due to the impaired recycling. The inhibited recycling of reduced lipoyl to oxidized lipoyl thus hinders the function of the enzymes. Collectively, we have further expanded the therapeutic scope of the bacterial salvage pathway to treat lipoylation deficiencies caused by defects in fatty acid synthesis (MECR) and lipoyl insertion accessory proteins (FDX1), but not by defects in lipoyl recycling (DLD).

**Figure 5.**
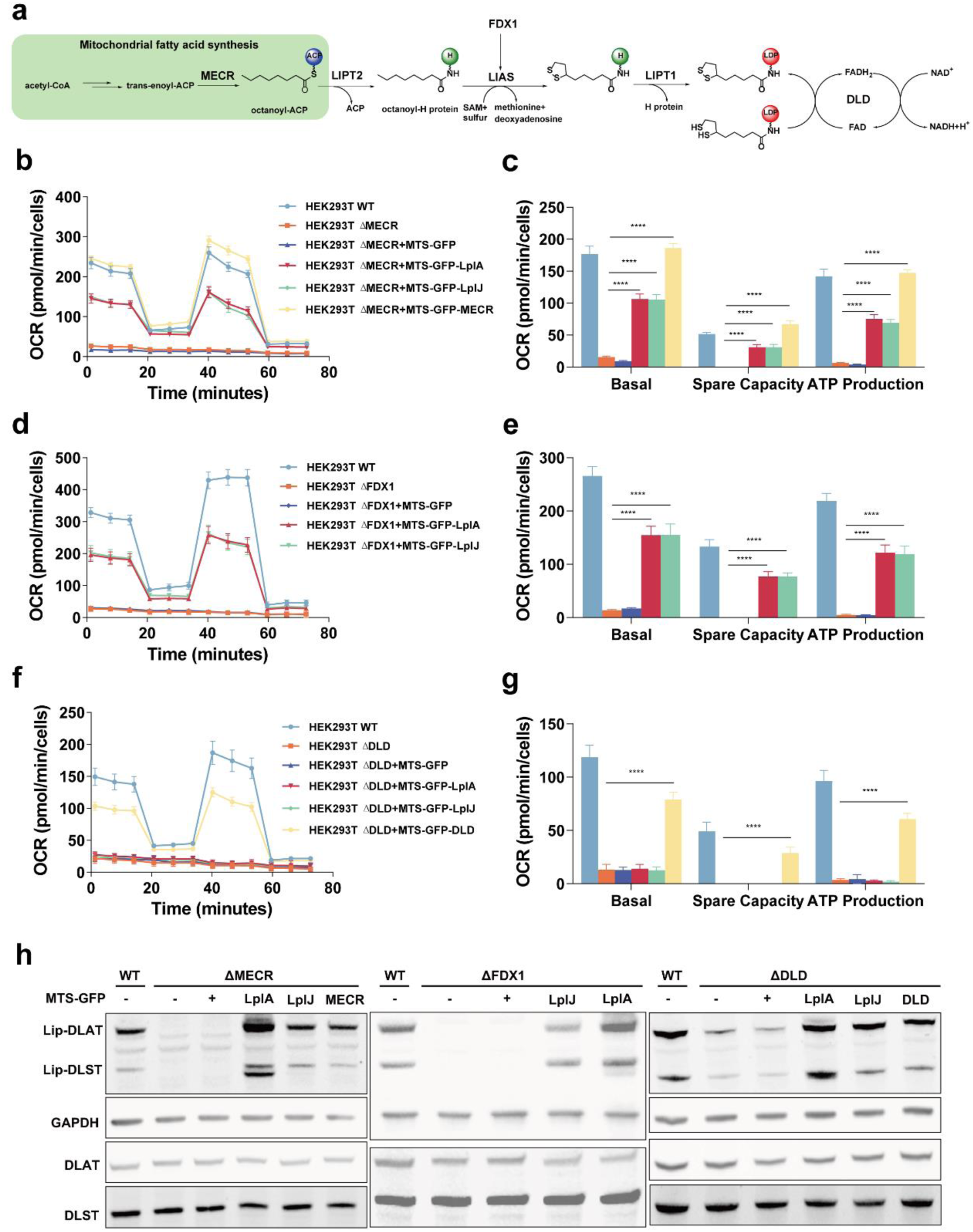
Expanding the therapeutic scope of LplA and LplJ. (a) Biosynthetic roles of MECR, FDX1 and DLD in human lipoylation pathway. (b) Mitochondrial respiration of HEK293T WT, ΔMECR and related complementary cells with lipoic acid supplementation. (c) OCR-related basal respiration, spare capacity and ATP production of ΔMECR related cells. (d) Mitochondrial respiration of HEK293T WT, ΔFDX1 and related complementary cells with lipoic acid supplementation. (e) OCR-related basal respiration, spare capacity and ATP production of ΔFDX1 related cells. (f) Mitochondrial respiration of HEK293T WT, ΔDLD and related complementary cells with lipoic acid supplementation. (g) OCR-related basal respiration, spare capacity and ATP production of ΔDLD related cells. (h) Lipoylation levels of DLAT and DLST of related cells were determined by Western blotting, GAPDH, total DLAT, and total DLST were used as controls. Data represented as mean±SD. Statistical significance was determined using multiple t tests (\**p* <0.05, \*\**p* <0.01, \*\*\**p* <0.001, and \*\*\*\**p* <0.0001).

### LplA ameliorates embryonic demise of homozygous LIPT1 knockout in mice

To evaluate the therapeutic potential of LplA *in vivo*, we used CRISPR-Cas9 to generate *Lipt1* knockout mice and *LplA* knockin mice. Interbreeding of *Lipt1*^+/-^ mice and Rosa26-LplA mice was performed to obtain *Lipt1*^+/-^ *LplA*^OE/+^ mice. Progeny carrying heterozygous *Lipt1*^+/-^ alleles were normal in growth and fecundity. Interbreeding between heterozygous *Lipt1*^+/-^ mice with oral administration of lipoic acid in water failed to give birth to any viable homozygous *Lipt1*^-/-^ mice (10 pups from 2 litters) (**Figure 6a**), which was consistent with previous study^25^. Excitingly, interbreeding between heterozygous *Lipt1*^+/-^ *LplA*^OE/+^ mice with oral administration of lipoic acid successfully yielded homozygous *Lipt1*^-/-^ mice carrying either heterozygous *LplA*^OE/+^ alleles or homozygous *LplA*^OE/OE^ alleles (9 pups out of 62 pups) (**Figure 6b**).

**Figure 6.**
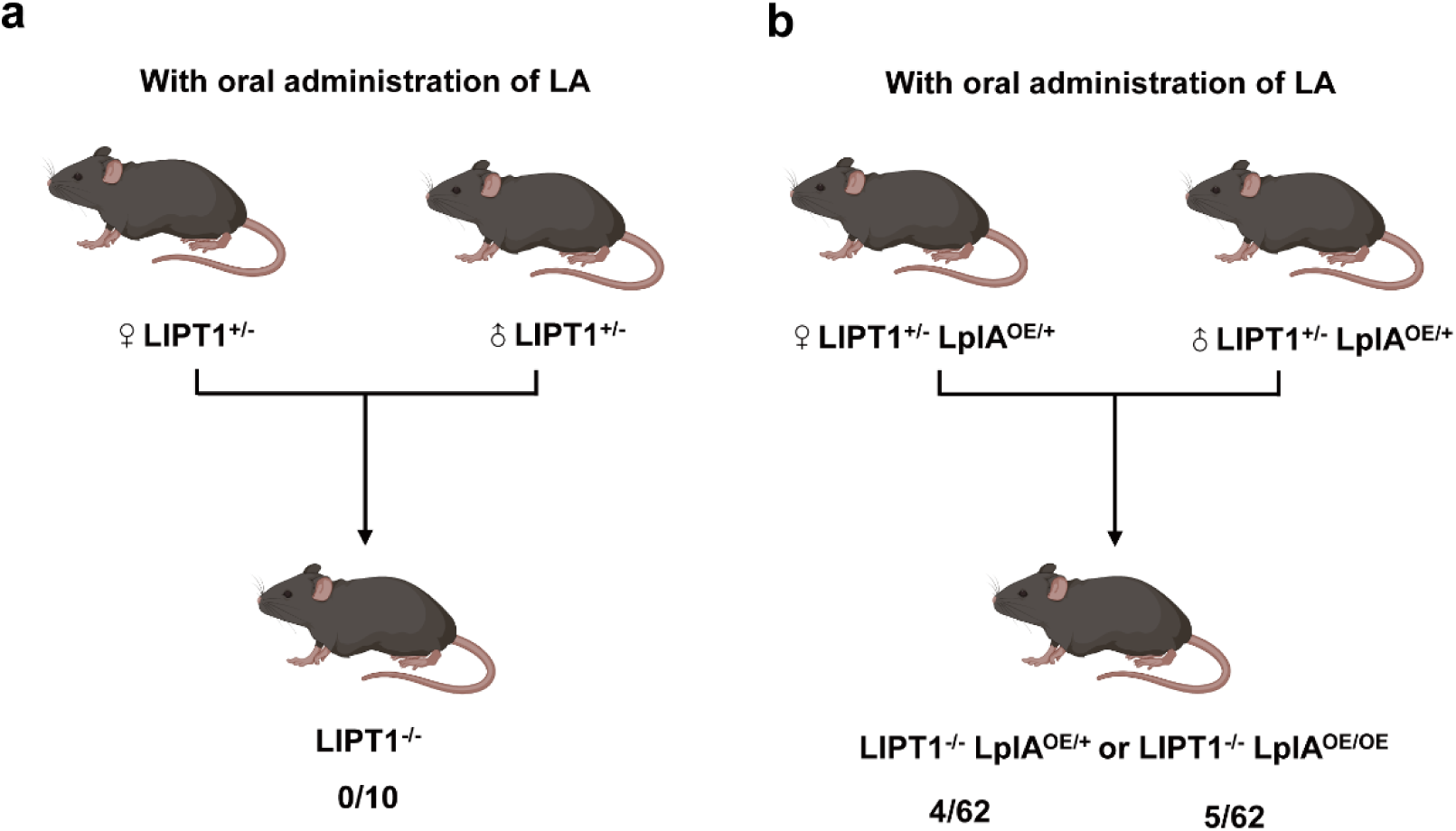
LplA ameliorates embryonic demise of homozygous LIPT1 knockout in mice. (a) Interbreeding strategy between heterozygous *Lipt1^+/-^* mice with oral administration of lipoic acid. The birth rate of *Lipt1^-/-^* mice is 0/10. (b) Interbreeding strategy between heterozygous *Lipt1^+/-^ LplA^OE/+^*mice with oral administration of lipoic acid. The birth rate of *Lipt1^-/-^*mice with *LplA^OE/+^* is 4/62. The birth rate of *Lipt1^-/-^*mice with *LplA^OE/OE^* is 5/62.

### *LplA^OE/+^* mice exhibit normal metabolic and immunological features

The introduction of bacterial proteins for therapeutic purposes raises concerns about safety. To further evaluate the safety of LplA treatment, we systematically examined the body composition, energy expenditure and hematology of *LplA^OE/+^* mice. EchoMRI analysis revealed that *LplA^OE/+^* mice had similar total body weight, fat mass and lean mass compared to wildtype mice (**Figure 7a**). Indirect calorimetry demonstrated that LplA expression increased whole-body oxygen consumption (VO_2_), carbon dioxide formation (VCO_2_), and energy expenditure throughout both dark and light cycles without altering respiration exchange ratio (**Figure 7b-c**). Hematological analysis revealed that the total white blood cell count (WBC), total red blood cell count (RBC), mean corpuscular volume (MCV), mean cellular hemoglobin content of erythrocytes (MCH), mean cellular hemoglobin concentration in erythrocytes (MCHC), red cell distribution width (RDW), platelet count (PLT), mean platelet volume (MPV), lymphocytes (Lymph) count, monocytes (Mono) count, neutrophil granulocytes (Neut) count, and eosinophil granulocytes (Eo) count in *LplA^OE/+^* mice exhibited normal levels to those of wildtype mice (**Figure 7d**). Protein lipoylation levels and PDH enzyme activities in muscle and heart of *LplA^OE/+^* mice showed no significant difference compared to that of wildtype mice (**Figure 7e-f**).

**Figure 7.**
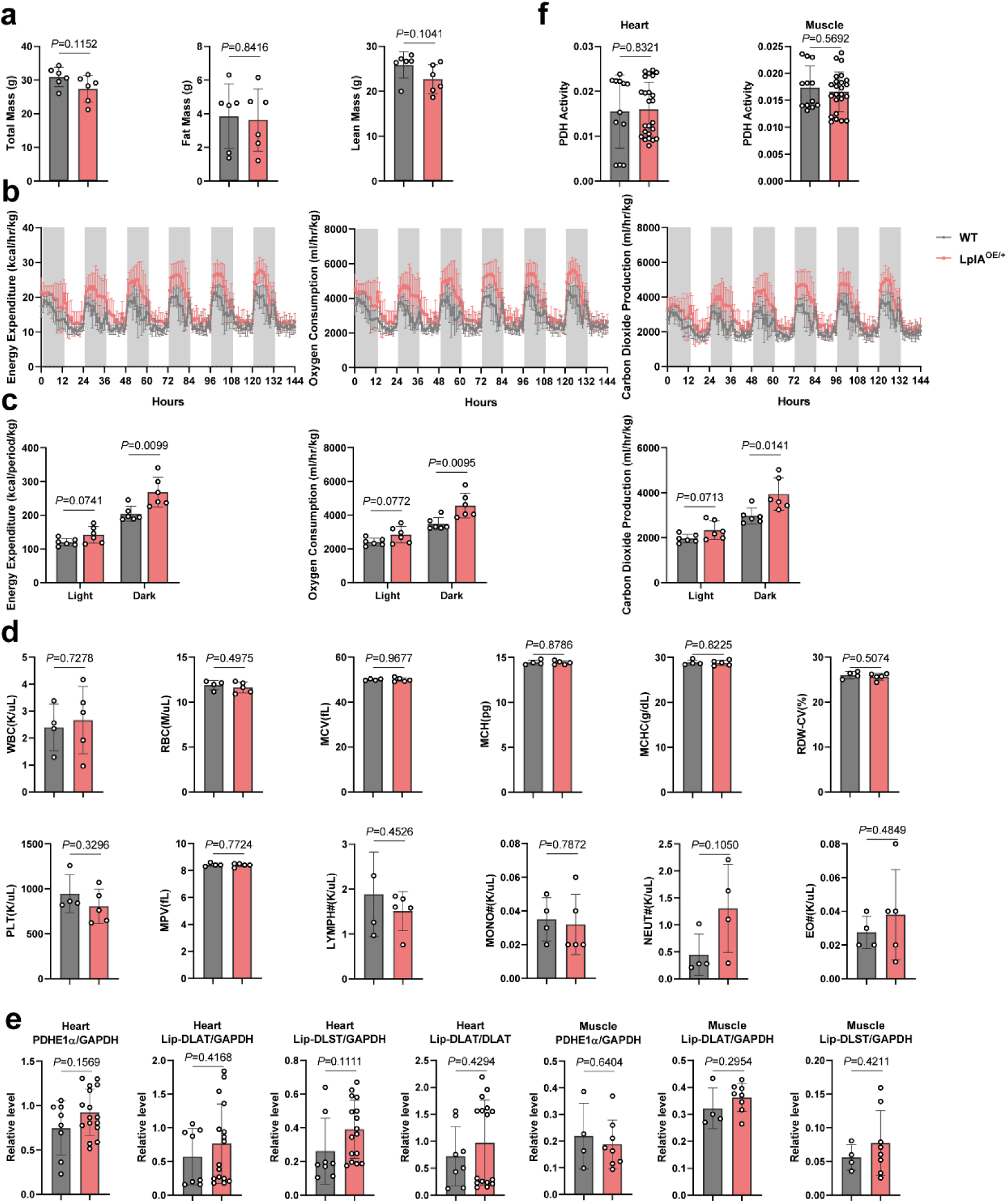
*LplA^OE/+^* mice exhibit normal metabolic and immunological features. (a) Total body weight, fat mass and lean mass of *LplA^OE/+^* mice and wildtype mice. (b) Light- and dark-cycle measurements of oxygen consumption (VO_2_), carbon dioxide output (VCO_2_), and energy expenditure of *LplA^OE/+^* mice and wildtype mice as quantified by indirect calorimetry. The dark shaded area indicates the dark cycle, and the clear open area indicates the light cycle. (c) Quantified VO_2_, VCO_2_, and energy expenditure during the light and dark cycle. (d) Total white blood cell count (WBC), total red blood cell count (RBC), mean corpuscular volume (MCV), mean cellular hemoglobin content of erythrocytes (MCH), mean cellular hemoglobin concentration in erythrocytes (MCHC), red cell distribution width (RDW), platelet count (PLT), mean platelet volume (MPV), lymphocytes (Lymph) count, monocytes (Mono) count, neutrophil granulocytes (Neut) count, and eosinophil granulocytes (Eo) count in *LplA^OE/+^* mice and wildtype mice. (e) The lipoylation level of DLAT and DLST in the heart and muscle in *LplA^OE/+^* mice and wildtype mice as quantified by Western blotting analysis. GAPDH was used as control. (f) PDH activities in the heart and muscle in *LplA^OE/+^* mice and wildtype mice. Data represented as mean±SD. Each data point represents an individual mouse. Exact *p* values are shown. Statistical significance was determined using multiple t tests.

## Conclusion

In this study, we validated the effectiveness and safety of bacterial lipoate protein ligases in rescuing inborn errors of metabolism caused by lipoylation deficiencies *in vitro* and *in vivo*. The robustness of this strategy lies in its potential to address disorders caused by defects in multiple genes, making it a highly appealing approach of gene therapy development for rare genetic diseases. More efforts are warranted to pave the way to clinical translation, including the optimization of gene delivery methods and the evaluation of long-term safety and efficacy.

---

**Figure S1.**
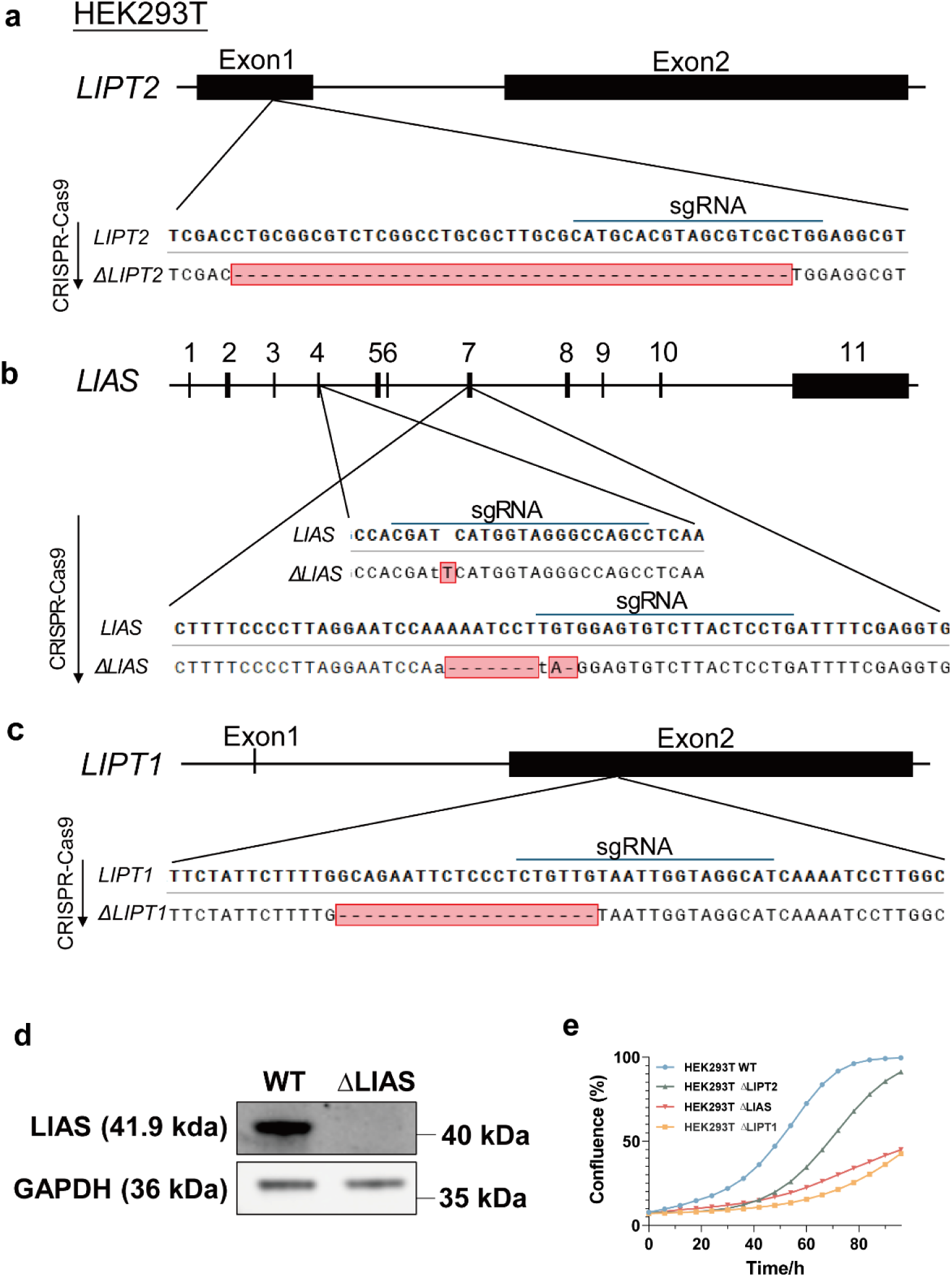
Characterization of the knockout cells. (a) LIPT2 knockout strategy through CRISPR-Cas9 in HEK293T cells and alignment of single colony sequencing. (b) LIAS knockout strategy through CRISPR-Cas9 in HEK293T cells and alignment of single colony sequencing. (c) LIPT1 knockout strategy through CRISPR-Cas9 in HEK293T cells and alignment of single colony sequencing. (d) Western blotting analysis of HEK293T WT and ΔLIAS cells using anti-LIAS antibody. LIAS band was abolished in ΔLIAS cells. (e) Cell growth curves of HEK293T WT, ΔLIPT2, ΔLIAS, and ΔLIPT1 cells with lipoic acid supplementation as determined by Incucyte live cell analysis system.

**Figure S2.**
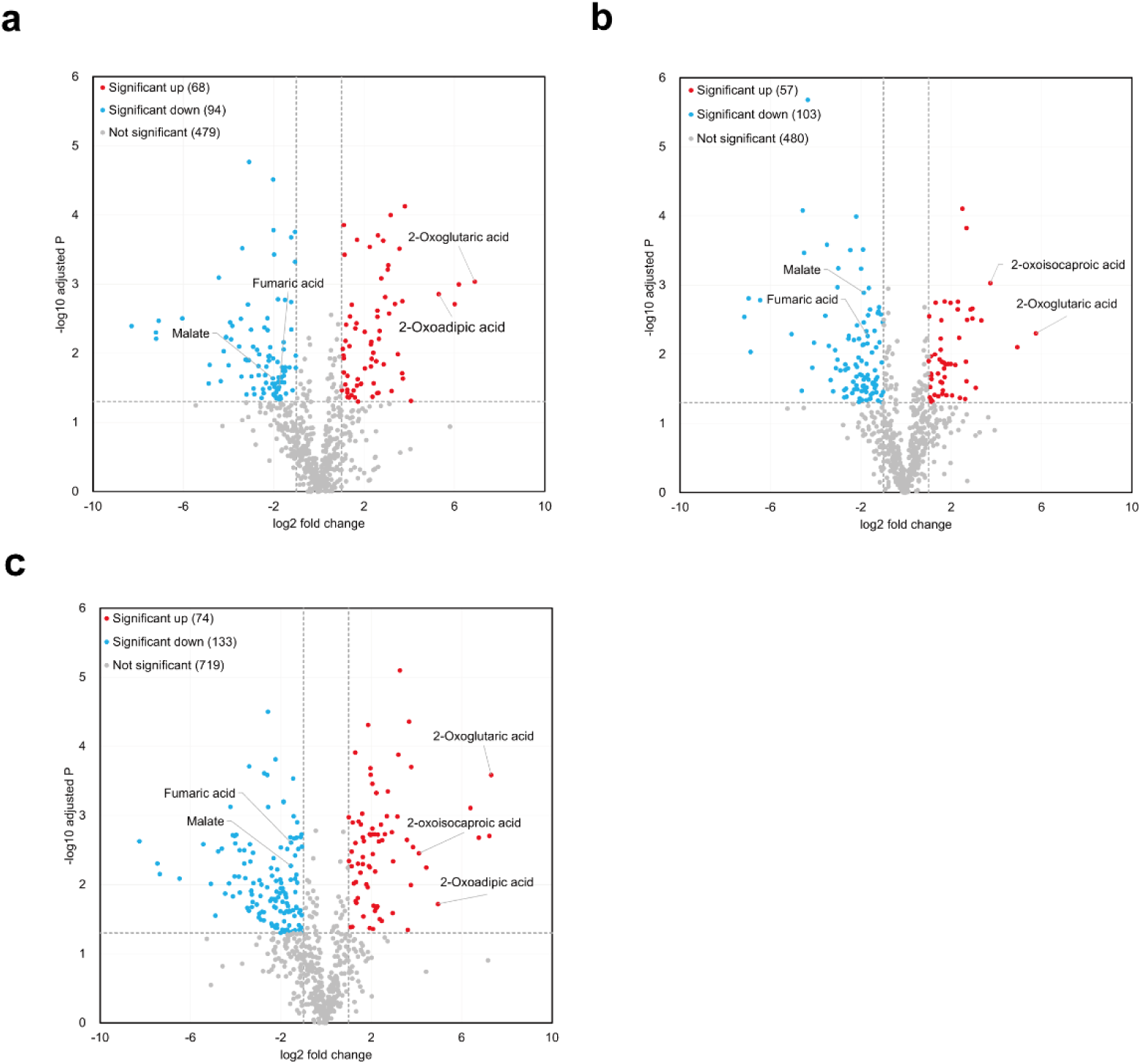
Untargeted metabolomics analysis of knockout cells compared to wildtype cells. (a) Volcano map of metabolomics of ΔLIPT2 compared to wildtype cells. (b) Volcano map of metabolomics of ΔLIAS compared to wildtype cells. (c) Volcano map of metabolomics of ΔLIPT1 compared to wildtype cells.

**Figure S3.**
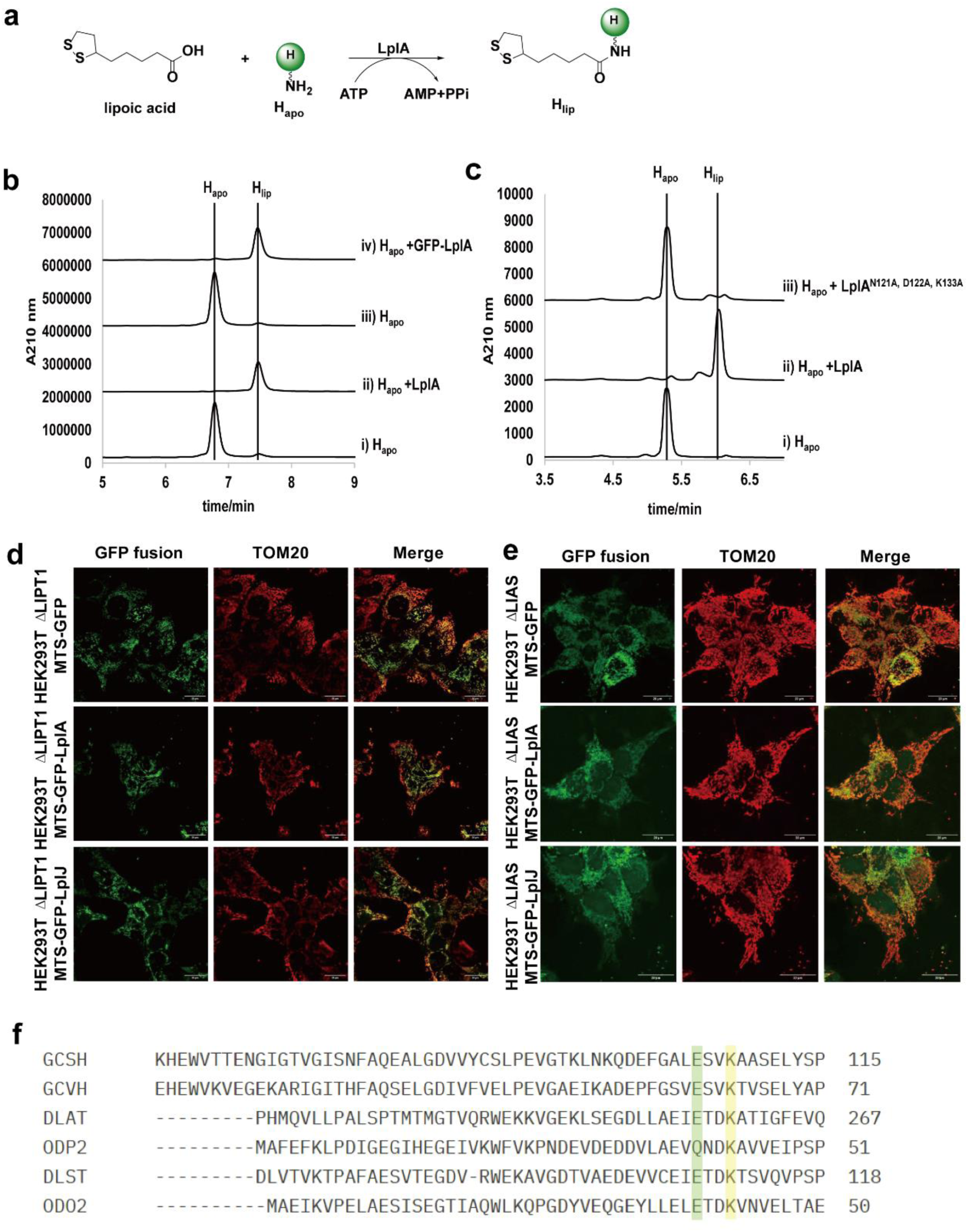
Cellular localization of complementary cells and lipoylation function characterization of LplA mutants. (a) Immunofluorescence analysis of HEK293T ΔLIPT1 MTS-GFP, MTS-GFP-LplA, and MTS-GFP-LplJ cells confirmed mitochondrial localization as monitored by confocal microscopy. Immunofluorescence analysis of HEK293T ΔLIAS MTS-GFP, MTS-GFP-LplA, and MTS-GFP-LplJ cells confirmed mitochondrial localization. (b) Scheme of lipoylation reaction of H protein catalyzed by LplA. (c) UV traces of lipoylation assay with LplA and GFP-LplA at 210 nm. (d) UV traces of lipoylation assay with LplA and LplA^N121A, D122A, K133A^ at 210 nm. (e) Amino acids sequence alignments of human and *B. subtilis*-derived E2 subunits and H protein. The conserved lysine modification site was marked yellow. The lipoylation recognition glutamate site was marked green. GCSH: human H protein, GCVH: *B. subtilis* H protein, DLAT: human PDH E2 subunit, ODP2: *B. subtilis* PDH E2 subunit, DLST: human OGDH E2 subunit, ODO2: *B. subtilis* OGDH E2 subunit.

**Figure S4.**
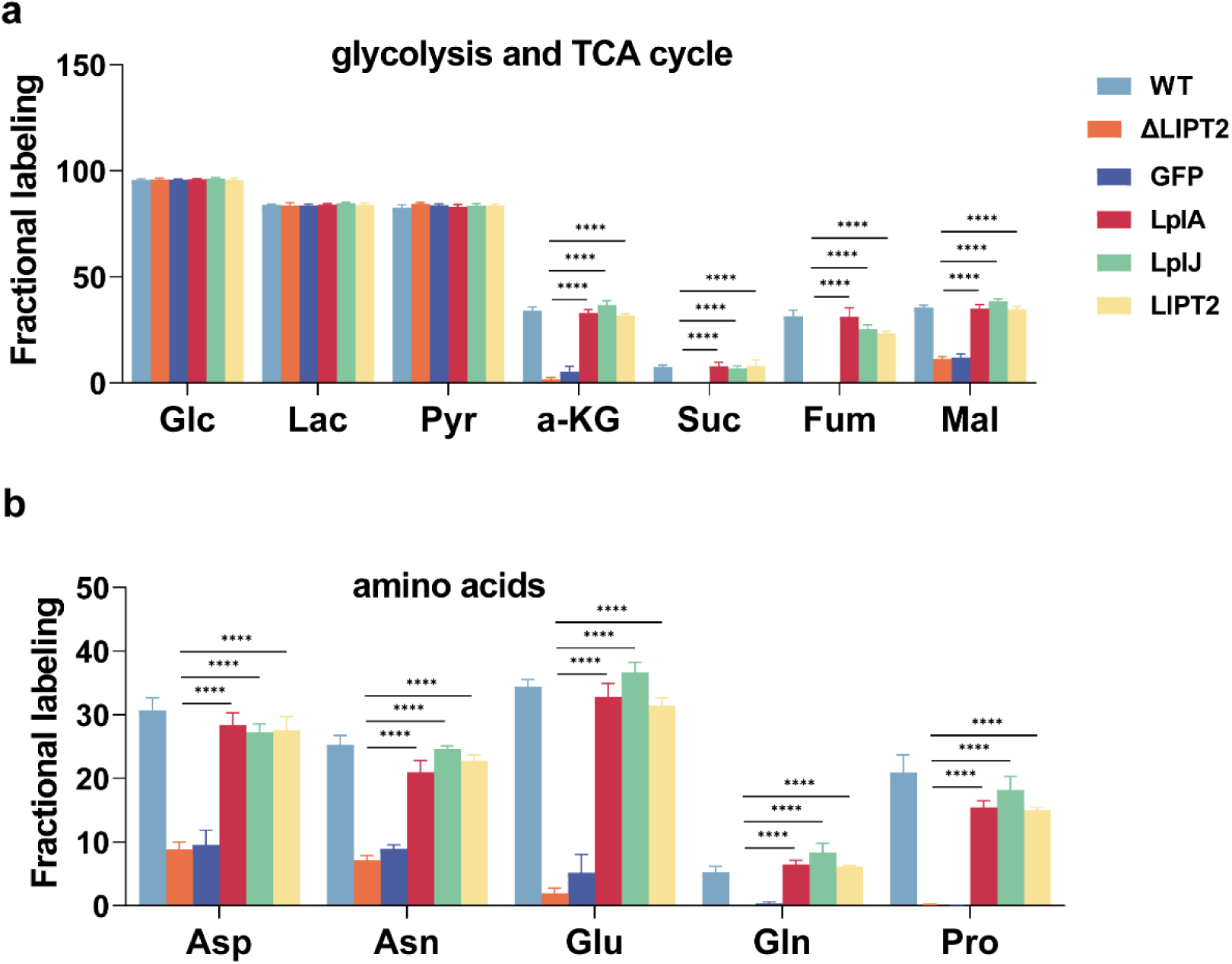
Isotope tracing analysis of HEK293T ΔLIPT2 and related complementary cells. (a) Fractional labeling of glycolysis and TCA cycle intermediates of HEK293T WT, ΔLIPT2, and ΔLIPT2 complemented with MTS-GFP, MTS-GFP-LplA, MTS-GFP-LplJ and MTS-GFP-LIPT2 cells after U-13C-glucose feeding. (b) Fractional labeling of amino acids intermediates of HEK293T WT, ΔLIPT2, and Δ LIPT2 complemented with MTS-GFP, MTS-GFP-LplA, MTS-GFP-LplJ and MTS-GFP-LIPT2 cells after U-13C-glucose feeding. Data represented as mean±SD. Statistical significance was determined using multiple t tests (\**p* <0.05, \*\**p* <0.01, \*\*\**p* <0.001, and \*\*\*\**p* <0.0001).

## Methods

### Key Resources Table

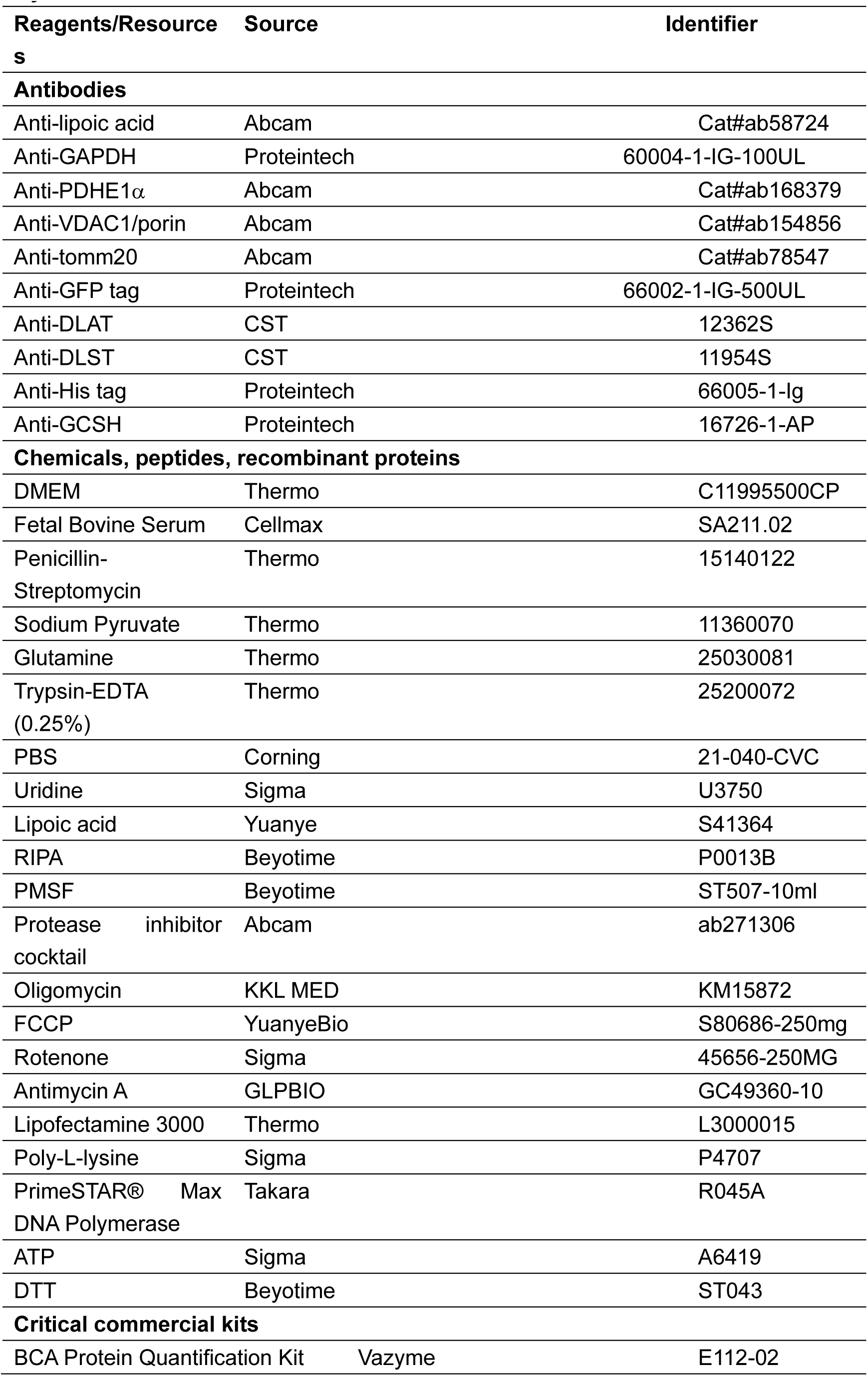
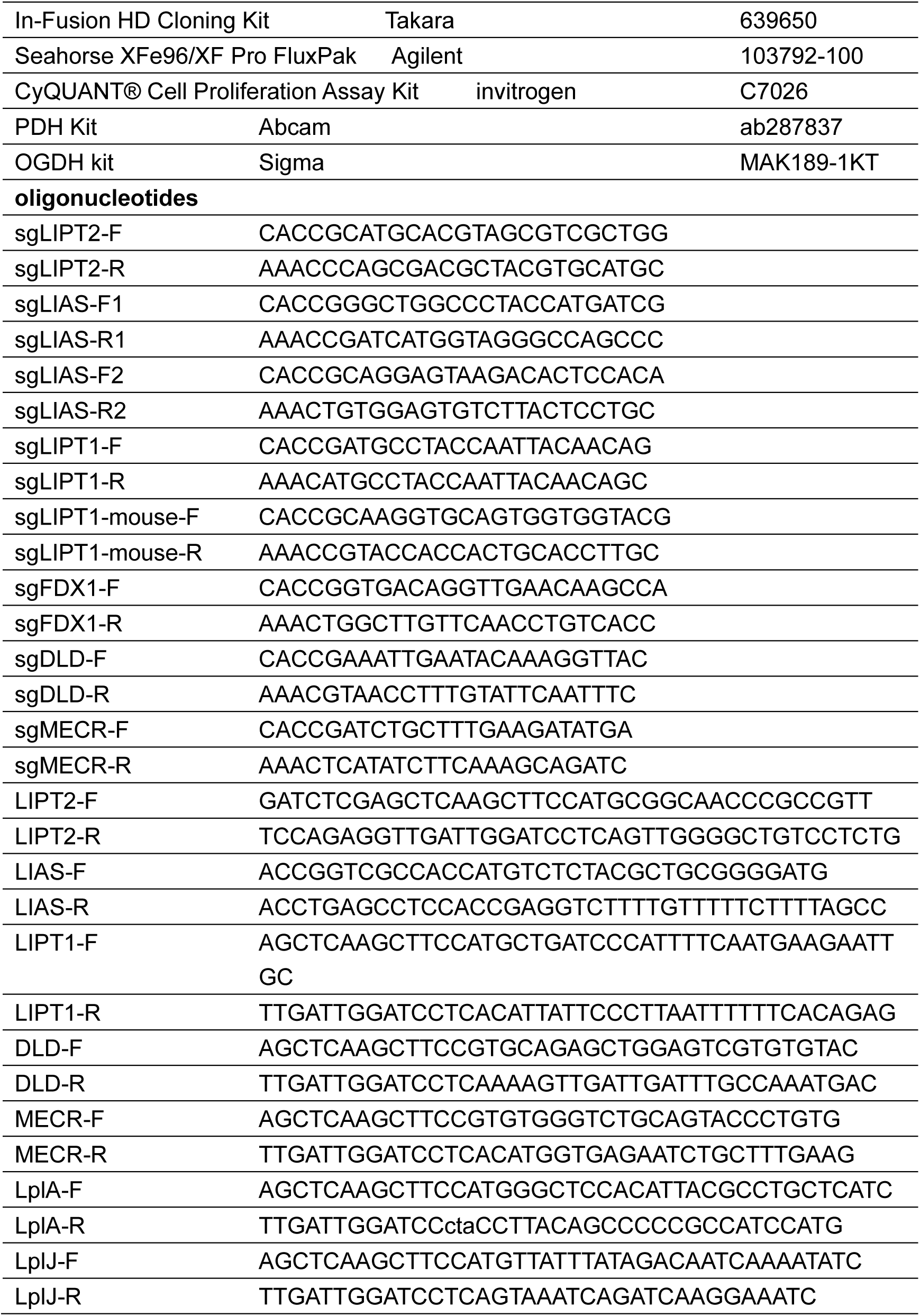

## Methods

### Cell Lines and Cell Culture Conditions

HEK293T cell line was obtained from the National Collection of Authenticated Cell Cultures (Serial No. SCSP-502). WT SV40 MEF cell line was obtained from ATCC (CRL-2907). The wildtype cells, gene knockout cells and related overexpressing stable cells were cultured in DMEM high glucose medium (Gibco, USA) supplemented with 10% FBS (CellMax), 100 U/mL penicillin-streptomycin, 50 μg/mL uridine and additional 1 mM pyruvate, and maintained in 5% CO_2_ incubator at 37°C.

### CRISPR-Cas9-Mediated Knockout Cells

sgRNAs for each gene (*LIPT2, LIAS, LIPT1, FDX1, DLD, MECR*) were designed through public website https://portals.broadinstitute.org/gppx/crispick/public and synthesized DNA oligos were ligated into BbsI-(NEB) digested px458 vector (Addgene, 48138)^37^. Targeting vectors were verified by Sanger sequencing. WT HEK293T or MEF cells were transiently transfected with px458-sgRNA vector for 3 days. Single cell clones were obtained by FITC-guided sorting in 96-well plates and successful knockout was verified by antibody staining and Hi-TOM sequencing. Briefly, a pair of primers containing bridging sequences at 5′ ends (5′-ggagtgagtacggtgtgc-3′ and 5′-gagttggatgctggatgg-3′) were used to amplify the sgRNA region at the first round of PCR amplification. Subsequently, barcoding PCR was conducted using the first round of PCR products as templates, followed by high-throughput sequencing and further decoding by a Hi-TOM platform (China National Rice Research Institute, Chinese Academy of Agricultural Sciences, Hangzhou)^38^.

### Constructs

DNA fragments encoding *E. coli* LplA and *B. subtilis* LplJ were PCR amplified from *E. coli* and *B. subtilis* genomic DNA respectively. DNA fragments encoding human *LIPT1*, *LIAS*, *LIPT2*, *DLD*, *MECR* were PCR amplified from U2OS or HEK293T cDNA. MTS sequence of ornithine transcarbamylase (OTC, mitochondrial) was synthesized by Genscript company (Nanjing, China). GFP fragment was PCR amplified from mEGFP-C1 plasmid (Addgene, 54759). MTS fragment, GFP Fragment and each gene fragment were inserted into pCDH-CMV (Addgene, 72265) backbone using Takara In-Fusion HD Cloning Kit. All constructs were confirmed by sequencing.

### Overexpression Stable Cell Lines

For lentivirus transfections, HEK293T WT cells were seeded at 5 × 10^5^ cells/well in 6-well plates for 24h. 1 μg of pCDH plasmid, 500 ng of psPAX2, and 500 ng of pMD2.G were used according to the manual of lipofectamine 3000 reagent (Thermo). After two days, the supernatants were collected and filtered through 0.22 μm filters. 1 mL of the flowthrough, 1 mL of new medium and 2 μL polybrene (Solarbio) were mixed and added to HEK293T KO cells plated in 6-well plates at 1 × 10^6^ cells/well. Cells expressing strong GFP signals were collected by MA900 sorter (SONY).

### Mitochondrial Stress Test

A Seahorse XFe96 extracellular flux analyzer (Agilent, USA) was used to measure oxygen consumption rate (OCR). HEK293T cells were seeded at 20000 cells/well and incubated for 24 h. Before assay, cells were equilibrated for 1 h in a non-CO_2_ incubator with XF Base medium supplemented with 10 mM glucose, 2 mM pyruvate and 4 mM L-glutamine. OCR were measured through sequential injection of 2 μM oligomycin, 0.5 μM carbonyl cyanide-4 (trifluoromethoxy) phenylhydrazone (FCCP), 1 μM rotenone and 1 μM antimycin A. Mitochondrial functions were indicated as basal respiration, spare respiratory capacity, proton leak and ATP production. Data was normalized using cell proliferation assay kit (Invitrogen, C7026).

### Cell Growth Curve

An Incucyte Real-Time Live Cell Analysis System (Sartorius, Germany) was used to determine cell growth curve. HEK293T cells were seeded at 2500 cells/200 μL/well in transparent 96-well plate and incubated in 5% CO_2_ Incucyte at 37°C. 10X objective, phase imaging mode and 6 hours scanning interval were set up. Images of living cells over time were automatically acquired. Cell confluence (%) was analyzed by phase contrast images.

### Western Blot

Cells were washed with ice-cold PBS three times and resuspended in RIPA buffer containing 1 mM PMSF. After lysis on ice for 30 min and centrifuge at 4°C for 30 min, the supernatant was collected for protein quantification using Pierce BCA protein assay. Residual supernatant was boiled with 2 × protein SDS PAGE loading buffer (Takara) at 98°C for 10 min. Total 30 ug protein were loaded on SDS-polyacrylamide gel electrophoresis (PAGE) gels and transferred onto nitrocellulose (NC) membrane. The NC membranes were blocked in 5% skimmed milk in tris-buffered saline with tween 20 (TBST) at room temperature for 1 h and incubated overnight at 4°C with following primary antibodies: anti-lipoic acid antibody (Abcam, Cat#ab58724, 1:1000), GAPDH antibody (Proteintech, Cat#60004-1-Ig, 1:2000). Anti-rabbit immunoglobulin G (IgG) (LI-COR, Cat#926-32211, 1:10000) and anti-mouse IgG (LI-COR, Cat#926-68070, 1:10000) were applied as secondary antibodies. ChemiDoc MP Imaging system (12003154, BioRad) was used for imaging.

### Isotope tracing analysis

The cells were plated at 10 cm cell plates and allowed to adhere for 24 h before labeling. After 24 h, cells were cultured in glucose-free DMEM supplemented with 10% FBS, 1% Penicillin-Streptomycin, 1 mM sodium pyruvate, 50 μg/uL uridine, 20 μM lipoic acid, 2 mM glutamine and 4g/L 13C-U-Glucose for 24 h before sample collection. The cells were gently washed with ice-cold 1X PBS and placed on dry ice for further processing. 4 mL of pre-chilled 80% Methanol was added to the cells rapidly, which were then placed at -80 °C for 15 min. Subsequently, the cells were then scraped with cell scrapers. The mixtures were transferred into tubes. 1 mL of 80% methanol was used to rinse the dish, and the rinse was combined with the previously collected mixtures. The combined methanol extracts were centrifuged at 4000g for 10 minutes at 4°C. The supernatant was collected, while the pellet was resuspended in 1 mL of pre-chilled methanol, followed by a second round of centrifugation under the same conditions. The supernatant from this step was combined with the previously collected supernatant. The pooled supernatants were dried under a gentle stream of nitrogen gas and stored at -80°C until further analysis. The extracts were dissolved in 200 μL acetonitrile/methanol/water (2:2:1) solution for HPLC-QTOF analysis. Intracellular metabolites were quantitated by the Agilent 1290 Infinity II UHPLC system tandem with 6545 Q/TOF-MS system. Chromatographic separation was achieved on an

ACQUITY UPLC BEH Amide column (100mm×2.1 mm, 1.7 μm). The mobile phase consisted of 15 mM ammonium acetate, 0.3% NH_3_·H_2_O in water (A) and 15 mM ammonium acetate, 0.3% NH_3_·H_2_O in 9:1 acetonitrile/water (B) at a flow rate of 0.3 mL/min. In detail, the column was eluted with 95% mobile phase B for 1 minute, followed by a linear gradient to 50% mobile-phase B over 8 min, held at 50% for 3 min, a linear gradient to 95% mobile phase B over 0.5 min, then 1.5 min at 95% mobile-phase B. The sample volume injected was 5 μL. MS data were acquired using electrospray ionization in both positive and negative ion mode over 50-1250 m/z. The parameters used were as follows: sheath gas temperature, 350℃; sheath gas flow, 11 L/min; capillary voltage, 4000 V; nozzle voltage, 1000 V; gas temperature, 350℃; nebulizer gas, 30 psi; drying gas flow rate, 8 L/min; fragmentor, 110 V; skimmer, 65 V. Raw data were processed using Profinder 10.0 (Agilent) for peak detection, alignment and integration.

### Protein expression and purification

The gene encoding *E. coli* LplA was inserted into pET-28a (+) vectors by In-fusion cloning. Site-directed mutations were accomplished according to the protocol provided in a Mut Express II Fast Mutagenesis kit V2 (Vazyme, Jiangsu, China). Proteins were expressed in *E. coli* BL21(DE3). Recombinant cells were incubated at 37°C in Luria-Bertani medium containing 50 μg/mL kanamycin until the OD_600_ reached about 0.6, and 0.2 mM Isopropyl-β-D-Thiogalactoside (IPTG) was added to induce protein expression for 18 h at 16°C. Cells were then harvested by centrifugation at 10,000×g, 4°C for 10 min. Cell pellets were resuspended in Buffer A (300 mM NaCl, 50 mM Tris-HCl, pH 7.5) with 5 mM imidazole, and lysed by high-pressure homogenization. The lysed samples were centrifuged, and the supernatants were loaded onto and purified by a Ni^2+^ chelating Sepharose Fast Flow column (Cytiva), using an ÄKTA purifier (Cytiva). The recovered proteins were tested for purity on 4-20% SDS-PAGE. The purified proteins were concentrated, and buffer exchanged into Buffer A, using Amicon Ultra filters. The purified proteins were flash-frozen in liquid nitrogen and stored at −80 °C.

### Lipoylation *in vitro* assay

The reaction system for enzyme activity determination is given in **Table 1**. The reaction was started with the addition of ATP, incubated at 30 °C overnight, and then terminated by heating for 90s in boiling water to completely denature and precipitate LplA. The amount of lipoylated H-protein in the mixture was determined using HPLC as described in a previous study^39^.

**Table 1.**
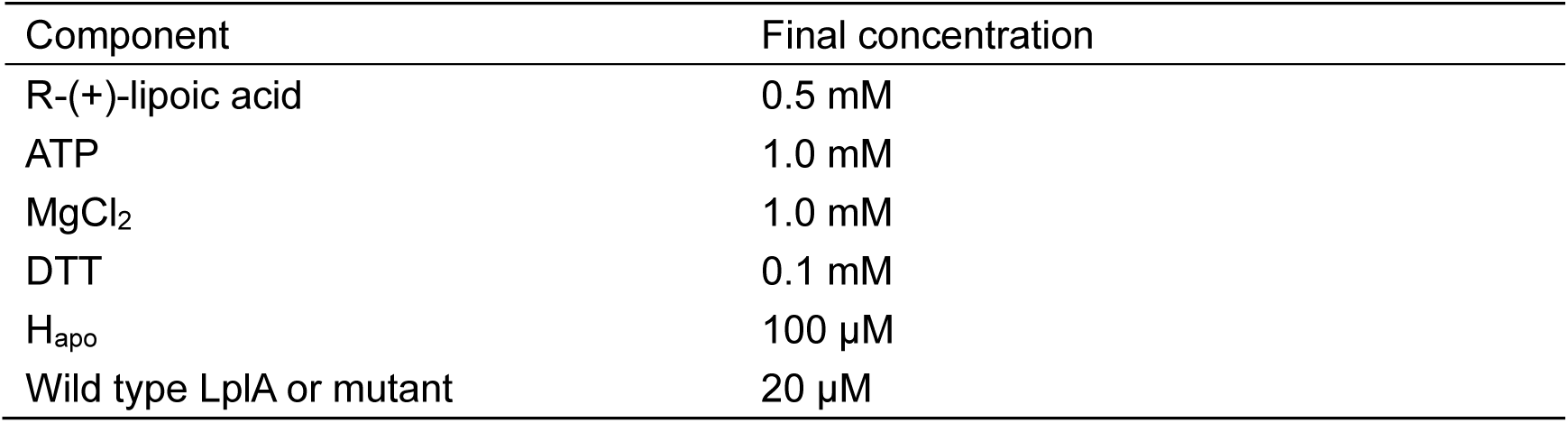
The reaction system for enzyme activity determination.

### Mice

All animal procedures were approved by the Institutional Animal Care and Use Committee (IACUC) at Westlake University and conducted in accordance with institutionally approved protocols and guidelines for animal care and use.

*Lipt1* knock-out mice and *LplA* knock-in mice were generated by the Laboratory Animal Resources Center at Westlake University using CRISPR/Cas9 genome editing technology.

C57BL/6J mice were used to generate founder mice. For *Lipt1* gene knockout, two sgRNAs were designed to delete most part of exon2, as exon2 included all coding region of *Lipt1*. Two specific allele sites were selected as target sequences in intron 1 and exon 2 of the mouse *Lipt1* locus. The two target sequences were sgRNA-2 (5-GAATCCAGAATCCTACTACC) and sgRNA-3 (5-GCCGTATGAGCAGGCGCATT).

Similarly, to generate KI mice expressing LplA, a construct containing the LplA cDNA was inserted into the mouse Rosa26 locus through homologous recombination using the CRISPR/Cas9 system. In brief, a CMV-OTC-LplA-polyA expression cassette was cloned into the *Roas26* construct and further verified by sequencing as targeting vector. The specific allele site was selected as target sequences (5-ACTCCAGTCTTTCTAGAAGA) between intron 1 and 2 of the mouse *Rosa26* locus.

The sgRNAs were produced by in vitro transcription (IVT). Briefly, sgRNAs were each synthesized from 1ug of plasmid DNA using a HiScribe T7 Quick High Yield RNA Synthesis Kitm kit (catalog number E2050; NEB) according to the manufacturer’s instructions. Each sgRNA was diluted to about 100ng/ul in RNase-free water. The sgRNAs and Cas9-NLS protein (catalog number M0646T; NEB) were mixed 1:1 to a final mixture for microinjection. For *LplA* KI mice generation, linearized targeting vector (5-10ng/ul) will be added into Cas9/sgRNA mix for pronuclear microinjection, the linearized DNA was purified using DNA Clean & Concentrator^®^-25 kit (Zymo research) and eluted in RNase-free water.

Cytoplasmic/pronuclear was performed in single-cell embryos derived from mating male and female C57BL/6 mice (stock number 000664; Jackson Laboratory, Bar Harbor, ME). Embryos were transferred into pseudo-pregnant B6/CBA F1 females, and resultant pups were screened for mutations by Sanger sequencing of PCR products (KO mice: forward primer, 5-AGTGAAAGCCAGAATGGCAG; reverse primer, 5-TATCCCCCTGATTTTTTCAC, target fragment less than 500bp). Mouse line heterozygous for *lipt1*-null alleles was established and designed *lipt1*^+/em38^. C57BL/6J WT mice were crossed to *lipt1*^+/em38^ founder mouse to confirm germline transmission.

Genotypes of KI mice were determined by amplifying a 1.9 kb product from purified mouse tail DNA (Rosa26 forward primer: 5-AAGGGAGCTGCAGTGGAGTA; Rosa26 reverse primer: 5-CCGAAAATCTGTGGGAAGTC). Mouse line heterozygous *Gt(ROSA)26Sor^+/em13^*^(^*^CMV-OTC-LplA^*^)^ was established and designed. C57BL/6J WT mouse were crossed to *LplA* founder mouse for germline transmission.

Interbreeding of *Lipt1*^+/-^ mice and Rosa26-LplA mice was performed to obtain *Lipt1*^+/-^; *LplA*^OE/+^ mice. Additionally, interbreeding of *Lipt1*^+/-^; *LplA*^OE/-^ male and female littermates was conducted to obtain *Lipt1*^-/-^; *LplA*^OE/+^ and *Lipt1*^-/-^; *LplA*^OE/OE^ mice.

### Genotyping method

Extraction of mice DNA using alkaline lysis method. Mouse toes or tails were lysed in 90 ul 50 mM sodium hydroxide at 95 °C for 1 h, followed by neutralization with 10 µl of 1 M Tris-HCl (pH 7.5). The lysate was then centrifuged at 12,000 ×g for 5 minutes. The supernatant which contained DNA was taken from the top of the tail prep avoiding the debris at the bottom of the well. The DNA was used as a template in 20 µl PCR reactions to amplify target sequences. The PCR products were analyzed by Sanger sequencing.

### PDH activity

100-200 mg minced mouse muscle tissue samples were cut into pieces and then homogenized with 1 mL ice-cold 1× Mitochondrial storage buffer (210 mM mannitol, 70 mM Sucrose, 5 mM Tris-HCl pH 7.5, 1 mM EDTA pH 7.5) using glass homogenizer on ice. The mixture was transferred to a fresh tube. 500 µl of 1×MS was used to wash the mortar. The wash buffer was also transferred to the tube. The homogenization buffer was centrifuged four times at 1,300 ×g for 10 min. Afterwards, the supernatant was transferred to a fresh tube and centrifuged at 10,000 ×g for 10 min. The pellet was collected for the following PDH assay.

Pyruvate dehydrogenase activity of mouse hind limb muscles was determined in vitro using PDH Kit (ab287837, Abcam) following the manufacturer’s instructions. Homogenized tissue was resuspended in 100 µl ice-cold PDH assay buffer and kept on ice for 10 min. 200 µl saturated ammonium sulfate were then added to lysate and keep on ice for 20 min to remove small molecules. After centrifuged at 10,000 ×g for 5 min, 100 µl PDH assay buffer was used to resuspend the pellet. The concentration was measured by Detergent Compatible Bradford Protein Quantification Kit (E211-01, Vazyme). The volume was then adjusted to 50 µl with appropriate concentration and mixed with 50 µl reaction buffer provided by PDH Kit. For background correction, 50 µl of background control mix was added to sample background control wells and mixed well. Absorbance was analyzed by Tecan Spark multimode microplate reader (Tencan Spark) at 450 nm in kinetic mode for 10-60 min, at 37 °C.

### Metabolic cages

Sixteen mice of six weeks, weighing approximately 20 g on arrival, were individually housed in Tecniplast 1284 cages (Tecniplast) for 6 days and acclimatized to standard animal house kept conditions. Temperature was kept at 23 °C with 40–70% humidity and lights were set to a 12 h:12 h light cycle with lights coming on at 7:00. Food pellets (Xietong Pharma Bio-eng. Co., Ltd.) and a chlorine residual of 1 to 2 ppm RO water were provided ad libitum throughout the entire experiment. The data were collected by the Promethion High-Definition Multiplexed Respirometry System (Sable Systems). The raw data was analyzed by online software (https://calrapp.org).

### Blood biochemistry test

Blood was collected from the mice orbital sinus. The eyeballs were quickly removed from the socket with a pair of tissue forceps. The body of the mouse was massaged so that blood was allowed to flow by capillary action into the lithium heparin tube (NPLH0505, SANLI). The samples were inverted gently for 30 seconds to mix. Within 30 minutes of collection, the samples were centrifuged at 12,000 xg for 5-10 min. Immediately after centrifugation, 300 μL of sample was transferred to a Catalyst* sample cup. The sample cup, slides, pipette tips were loaded in the sample drawer. The samples were analyzed by Catalyst One* Chemistry Analyzer (INDEXX).

### Complete Blood Count

Retro-orbital sampling is used to collect blood from mice. 3-4 drops of blood were collected with EDTAK_2_ tube (NPK230505, SANLI). The samples were inverted gently 10 times to mix. The samples were analyzed by the IDEXX ProCyte Dx® Hematology Analyzer (INDEXX).

## References

1. Rowland, E.A., Snowden, C.K., and Cristea, I.M. (2018). Protein lipoylation: an evolutionarily conserved metabolic regulator of health and disease. Current Opinion in Chemical Biology 42, 76–85. 10.1016/j.cbpa.2017.11.003.

2. Mayr, J.A., Feichtinger, R.G., Tort, F., Ribes, A., and Sperl, W. (2014). Lipoic acid biosynthesis defects. J of Inher Metab Disea 37, 553–563. 10.1007/s10545-014-9705-8.

3. Tort, F., Ferrer-Cortes, X., and Ribes, A. (2016). Differential diagnosis of lipoic acid synthesis defects. J of Inher Metab Disea 39, 781–793. 10.1007/s10545-016-9975-4.

4. Cronan, J.E. (2020). Progress in the Enzymology of the Mitochondrial Diseases of Lipoic Acid Requiring Enzymes. Front. Genet. 11, 510. 10.3389/fgene.2020.00510.

5. Hiltunen, J.K., Autio, K.J., Schonauer, M.S., Kursu, V.A.S., Dieckmann, C.L., and Kastaniotis, A.J. (2010). Mitochondrial fatty acid synthesis and respiration. Biochimica et Biophysica Acta (BBA) - Bioenergetics 1797, 1195–1202. 10.1016/j.bbabio.2010.03.006.

6. Cronan, J.E. (2024). Lipoic acid attachment to proteins: stimulating new developments. Microbiol Mol Biol Rev 88, e00005–24. 10.1128/mmbr.00005-24.

7. Mathias, R.A., Greco, T.M., Oberstein, A., Budayeva, H.G., Chakrabarti, R., Rowland, E.A., Kang, Y., Shenk, T., and Cristea, I.M. (2014). Sirtuin 4 Is a Lipoamidase Regulating Pyruvate Dehydrogenase Complex Activity. Cell 159, 1615–1625. 10.1016/j.cell.2014.11.046.

8. Cao, X., and Cronan, J.E. (2015). The Streptomyces coelicolor Lipoate-protein Ligase Is a Circularly Permuted Version of the Escherichia coli Enzyme Composed of Discrete Interacting Domains. Journal of Biological Chemistry 290, 7280–7290. 10.1074/jbc.M114.626879.

9. Hermes, F.A., and Cronan, J.E. (2013). The role of the *Saccharomyces cerevisiae* lipoate protein ligase homologue, Lip3, in lipoic acid synthesis. Yeast 30, 415–427. 10.1002/yea.2979.

10. Cao, X., Zhu, L., Song, X., Hu, Z., and Cronan, J.E. (2018). Protein moonlighting elucidates the essential human pathway catalyzing lipoic acid assembly on its cognate enzymes. Proc. Natl. Acad. Sci. U.S.A. 115. 10.1073/pnas.1805862115.

11. Cronan, J.E. (2016). Assembly of Lipoic Acid on Its Cognate Enzymes: an Extraordinary and Essential Biosynthetic Pathway. Microbiol Mol Biol Rev 80, 429–450. 10.1128/MMBR.00073-15.

12. Morris, T.W., Reed, K.E., and Cronan, J.E. (1994). Identification of the gene encoding lipoate-protein ligase A of Escherichia coli. Molecular cloning and characterization of the lplA gene and gene product. The Journal of Biological Chemistry 269, 16091–16100.

13. 13. A novel two-gene requirement for the octanoyltransfer reaction of Bacillus subtilis lipoic acid biosynthesis (2011). Molecular Microbiology 80, 335–349. doi:10.1111/j.1365-2958.2011.07597.x.

14. Paredes, F., Sheldon, K., Lassègue, B., Williams, H.C., Faidley, E.A., Benavides, G.A., Torres, G., Sanhueza-Olivares, F., Yeligar, S.M., Griendling, K.K., et al. (2018). Poldip2 is an oxygen-sensitive protein that controls PDH and αKGDH lipoylation and activation to support metabolic adaptation in hypoxia and cancer. Proc. Natl. Acad. Sci. U.S.A. 115, 1789–1794. 10.1073/pnas.1720693115.

15. Fujiwara, K., Takeuchi, S., Okamura-Ikeda, K., and Motokawa, Y. (2001). Purification, Characterization, and cDNA Cloning of Lipoate-activating Enzyme from Bovine Liver. Journal of Biological Chemistry 276, 28819–28823. 10.1074/jbc.M101748200.

16. Mayr, J.A., Zimmermann, F.A., Fauth, C., Bergheim, C., Meierhofer, D., Radmayr, D., Zschocke, J., Koch, J., and Sperl, W. (2011). Lipoic Acid Synthetase Deficiency Causes Neonatal-Onset Epilepsy, Defective Mitochondrial Energy Metabolism, and Glycine Elevation. The American Journal of Human Genetics 89, 792–797. 10.1016/j.ajhg.2011.11.011.

17. Tsurusaki, Y., Tanaka, R., Shimada, S., Shimojima, K., Shiina, M., Nakashima, M., Saitsu, H., Miyake, N., Ogata, K., Yamamoto, T., et al. (2015). Novel compound heterozygous LIAS mutations cause glycine encephalopathy. J Hum Genet 60, 631–635. 10.1038/jhg.2015.72.

18. Baker, P.R., Friederich, M.W., Swanson, M.A., Shaikh, T., Bhattacharya, K., Scharer, G.H., Aicher, J., Creadon-Swindell, G., Geiger, E., MacLean, K.N., et al. (2014). Variant non ketotic hyperglycinemia is caused by mutations in LIAS, BOLA3 and the novel gene GLRX5. Brain 137, 366–379. 10.1093/brain/awt328.

19. Wongkittichote, P. (2023). Novel LIAS variants in a patient with epilepsy and profound developmental disabilities. Molecular Genetics and Metabolism. Molecular Genetics and Metabolism 138, 107373. 10.1016/j.ymgme.2023.107373

20. Habarou, F., Hamel, Y., Haack, T.B., Feichtinger, R.G., Lebigot, E., Marquardt, I., Busiah, K., Laroche, C., Madrange, M., Grisel, C., et al. (2017). Biallelic Mutations in LIPT2 Cause a Mitochondrial Lipoylation Defect Associated with Severe Neonatal Encephalopathy. The American Journal of Human Genetics 101, 283–290. 10.1016/j.ajhg.2017.07.001.

21. Soreze, Y., Boutron, A., Habarou, F., Barnerias, C., Nonnenmacher, L., Delpech, H., Mamoune, A., Chrétien, D., Hubert, L., Bole-Feysot, C., et al. (2013). Mutations in human lipoyltransferase gene LIPT1cause a Leigh disease with secondary deficiency for pyruvate and alpha-ketoglutarate dehydrogenase. Orphanet J Rare Dis 8, 192. 10.1186/1750-1172-8-192.

22. Tort, F., Ferrer-Cortès, X., Thió, M., Navarro-Sastre, A., Matalonga, L., Quintana, E., Bujan, N., Arias, A., García-Villoria, J., Acquaviva, C., et al. (2014). Mutations in the lipoyltransferase *LIPT1* gene cause a fatal disease associated with a specific lipoylation defect of the 2-ketoacid dehydrogenase complexes. Human Molecular Genetics 23, 1907–1915. 10.1093/hmg/ddt585.

23. Taché, V., Bivina, L., White, S., Gregg, J., Deignan, J., Boyadjievd, S.A., and Poulain, F.R. (2016). Lipoyltransferase 1 Gene Defect Resulting in Fatal Lactic Acidosis in Two Siblings. Case Reports in Obstetrics and Gynecology. Case Reports in Obstetrics and Gynecology. 10.1155/2016/6520148

24. Stowe, R.C., Sun, Q., Elsea, S.H., and Scaglia, F. (2018). LIPT1 deficiency presenting as early infantile epileptic encephalopathy, Leigh disease, and secondary pyruvate dehydrogenase complex deficiency. American J of Med Genetics Pt A 176, 1184–1189. 10.1002/ajmg.a.38654.

25. Ni, M., Solmonson, A., Pan, C., Yang, C., Li, D., Notzon, A., Cai, L., Guevara, G., Zacharias, L.G., Faubert, B., et al. (2019). Functional Assessment of Lipoyltransferase-1 Deficiency in Cells, Mice, and Humans. Cell Reports 27, 1376–1386.e6. 10.1016/j.celrep.2019.04.005.

26. Feng, D., Witkowski, A., and Smith, S. (2009). Down-regulation of Mitochondrial Acyl Carrier Protein in Mammalian Cells Compromises Protein Lipoylation and Respiratory Complex I and Results in Cell Death. Journal of Biological Chemistry 284, 11436–11445. 10.1074/jbc.M806991200.

27. Liu, Z., Shimura, M., Zhang, L., Zhang, W., Wang, J., Ogawa-Tominaga, M., Wang, J., Wang, X., Lv, J., Shi, W., et al. (2021). Whole exome sequencing identifies a novel homozygous MECR mutation in a Chinese patient with childhood-onset dystonia and basal ganglia abnormalities, without optic atrophy. Mitochondrion 57, 222–229. 10.1016/j.mito.2020.12.014.

28. Ambrus, A., and Adam-Vizi, V. (2013). Molecular dynamics study of the structural basis of dysfunction and the modulation of reactive oxygen species generation by pathogenic mutants of human dihydrolipoamide dehydrogenase. Archives of Biochemistry and Biophysics 538, 145–155. 10.1016/j.abb.2013.08.015.

29. Gómez-Fernández, D., Romero-González, A., Suárez-Rivero, J.M., Cilleros-Holgado, P., Álvarez-Córdoba, M., Piñero-Pérez, R., Romero-Domínguez, J.M., Reche-López, D., López-Cabrera, A., Ibáñez-Mico, S., et al. (2024). A Multi-Target Pharmacological Correction of a Lipoyltransferase LIPT1 Gene Mutation in Patient-Derived Cellular Models. Antioxidants 13, 1023. 10.3390/antiox13081023.

30. Pietikäinen, L.P., Rahman, M.T., Hiltunen, J.K., Dieckmann, C.L., and Kastaniotis, A.J. (2021). Genetic dissection of the mitochondrial lipoylation pathway in yeast. BMC Biol 19, 14. 10.1186/s12915-021-00951-3.

31. Rasetto, N.B., Lavatelli, A., Martin, N., and Mansilla, M.C. (2019). Unravelling the lipoyl-relay of exogenous lipoate utilization in *Bacillus subtilis*. Molecular Microbiology 112, 302–316. 10.1111/mmi.14271.

32. Fujiwara, K., Okamura-Ikeda, K., and Motokawa, Y. (1997). [20] Lipoate addition to acyltransferases of α-keto acid dehydrogenase complexes and H-protein of glycine cleavage system. In Methods in Enzymology (Elsevier), pp. 184–193. 10.1016/S0076-6879(97)79022-0.

33. Craven, L., Alston, C.L., Taylor, R.W., and Turnbull, D.M. (2017). Recent Advances in Mitochondrial Disease. Annu. Rev. Genom. Hum. Genet. 18, 257–275. 10.1146/annurev-genom-091416-035426.

34. Dreishpoon, M.B., Bick, N.R., Petrova, B., Warui, D.M., Cameron, A., Booker, S.J., Kanarek, N., Golub, T.R., and Tsvetkov, P. (2023). FDX1 regulates cellular protein lipoylation through direct binding to LIAS. Journal of Biological Chemistry 299, 105046. 10.1016/j.jbc.2023.105046.

35. Joshi, P.R., Sadre, S., Guo, X.A., McCoy, J.G., and Mootha, V.K. (2023). Lipoylation is dependent on the ferredoxin FDX1 and dispensable under hypoxia in human cells. Journal of Biological Chemistry 299, 105075. 10.1016/j.jbc.2023.105075.

36. Schulz, V., Basu, S., Freibert, S.-A., Webert, H., Boss, L., Mühlenhoff, U., Pierrel, F., Essen, L.-O., Warui, D.M., Booker, S.J., et al. (2023). Functional spectrum and specificity of mitochondrial ferredoxins FDX1 and FDX2. Nat Chem Biol 19, 206–217. 10.1038/s41589-022-01159-4.

37. Ran, F.A., Hsu, P.D., Wright, J., Agarwala, V., Scott, D.A., and Zhang, F. (2013). Genome engineering using the CRISPR-Cas9 system. Nature Protocols 8, 2281–2308. DOI: 10.1038/nprot.2013.143.

38. Sun, T., Liu, Q., Chen, X., Hu, F., and Wang, K. (2024). Hi-TOM 2.0: an improved platform for high-throughput mutation detection. Sci. China Life Sci. 67, 1532–1534. 10.1007/s11427-024-2555-x.

39. Zhang, X., Nie, J., Zheng, Y., Ren, J., and Zeng, A. (2020). Activation and competition of lipoylation of H protein and its hydrolysis in a reaction cascade catalyzed by the multifunctional enzyme lipoate–protein ligase A. Biotech & Bioengineering 117, 3677– 3687. 10.1002/bit.27526.

